# Non-decameric NLRP3 forms an MTOC-independent inflammasome

**DOI:** 10.1101/2023.07.07.548075

**Authors:** María Mateo-Tórtola, Inga V. Hochheiser, Jelena Grga, Jana S. Mueller, Matthias Geyer, Alexander N.R. Weber, Ana Tapia-Abellán

## Abstract

NLRP3 is an inflammasome-forming protein that plays a key role in conditions ranging from infections to Alzheimer’s disease. NLRP3 is activated by many potassium (K^+^)-dependent or - independent stimuli related to pathogens or sterile insults. Molecularly, human NLRP3 assembles into a decameric ‘cage’ and its interaction with both the trans-Golgi network (TGN) and the microtubule organization centre (MTOC) has been proposed as critical for NLRP3 activation. However, the relative mechanistic requirement for K^+^-dependent versus - independent stimuli is unclear. Using spatially and dynamically high-resolution microscopy in human macrophages, we found that K^+^-dependent NLRP3 stimulation triggers two distinct activation pathways: (*i*) the decameric cage-dependent pathway, which was also exclusively required for the K^+^-independent stimulus, imiquimod; and (*ii*) a cage- and TGN/MTOC-independent pathway that was fully functional in an NLRP3 protein engineered to form monomers and unable to interact with membrane lipids. Collectively, our results delineate two parallel yet biologically distinct NLRP3 activation pathways.

## Introduction

Inflammation is an essential mechanism initiated by the innate immune system to quickly and effectively respond to threats. Inflammatory cytokines include members of the interleukin-1 (IL-1) family whose processing and release is regulated by cytosolic multiprotein complexes named inflammasomes, that are formed in inflammatory cells such as macrophages ^1^. Among them, NLRP3 is the most studied and strongly disease-associated inflammasome. *NLRP3* gain-of-function mutations are responsible for a range of genetic autoinflammatory diseases named Cryopyrin Associated Periodic Syndromes (CAPS) ^2, 3^. Furthermore, inappropriate activation of the NLRP3 inflammasome has been specifically associated with many physiological and pathological conditions in humans, from gout to Alzheimer’s disease ^4^. Thus, elucidating the mechanism of NLRP3 inflammasome activation has been the focus of intense research efforts ^5^.

The NLRP3 sensor is activated by a wide variety of stimuli related to microorganisms or sterile danger signals, and thus functions as a general sensor of cell homeostasis, whose minimum requirement to become active appears to be the intracellular loss of potassium (K^+^) ^6, 7^. Additionally, NLRP3 is activated by an K^+^-independent stimulus named imiquimod ^8^. However, the exact mechanism of how these stimuli activate NLRP3 is incompletely known. In general, upon activation, NLRP3 is thought to suffer a conformational change ^9, 10^ that allows the binding to the next protein of the pathway, the adaptor ASC. Then, ASC forms prion-like filaments that engage the effector protein, caspase-1, to form an active NLRP3 inflammasome. Active caspase-1 cleaves inflammatory cytokines such as pro-IL-1β to its active form and processes the pore-forming protein gasdermin D (GSDMD), thereby causing a type of inflammatory cell death known as pyroptosis and the release of mature IL-1β and other alarmins ^1, 11^.

Recently, our knowledge of the NLRP3 inflammasome has changed and increased in complexity by implicating NLRP3 oligomerization and its dynamic subcellular localization as critical determinants of the activation mechanism ^5, 12^. Cryo-EM studies showed that inactive NLRP3 forms steady-state oligomers resembling a ‘cage’. These cages consist of a decamer in humans and a dodecamer in mice ^13, 14^. Mutants presenting a disruptive cage were considered unable to form active NLRP3 inflammasomes in mouse cells ^13^. These results changed previous assumptions based e.g. on the NLRC4 inflammasome ^15, 16^, in which inactive NLRP3 was thought to be a monomer and only the active NLRP3 an oligomer ^17, 18^. Whereas smaller NLRP3 species such as monomers were obtained from cytosolic extracts of immortalized bone marrow-derived macrophages (iBMDM) and considered inactive ^13^, ectopic expression of an NLRP3 construct lacking the LRR, a domain critical for oligomer cage formation, still formed active inflammasomes, complicating the understanding of how non-oligomeric species can function ^19^. Recently, the best explored specific NLRP3 inhibitor, MCC950, was shown to directly bind NLRP3 in its central domain arresting the protein in the inactive conformation ^9, 13, 18^.

Regarding the role of subcellular localization in the activation pathway, NLRP3 was proposed to travel from the dispersed trans-Golgi network (TGN) to other membranes such as endosomes, until it reaches the microtubule organization centre (MTOC), where it somehow engages ASC and caspase-1 to form an active inflammasome ^3, 20–23^. Here, the affinity of a polybasic region in the inactive NLRP3 cage to bind negatively charged membrane lipids was shown to be critical for TGN recruitment and subsequent activation in murine NLRP3 ^13, 24^. However, the biological importance of the human NLRP3 decamer in the proposed TGN/MTOC activation pathway and the role of smaller non-oligomer/-cage NLRP3 species are not yet fully understood. Additionally, how NLRP3 structure, binding/release from membranes and trafficking interrelate, remains enigmatic.

By precisely deleting the complete exon 3 of *NLRP3*, we obtained an NLRP3 sensor lacking a short linker region and this protein was unable to organise in decamers and to bind lipids. When reconstituted in human THP-1 macrophages we observed that these smaller NLRP3 species were still capable of forming active inflammasomes, but completely independent and spatially distal of TGN and MTOC. Furthermore, finely time-resolved live cell microscopy showed that even NLRP3 wild-type (WT) could engage ASC in two different NLRP3 activation pathways, one MTOC-dependent and one MTOC-independent. Unexpectedly, K^+^-dependent NLRP3 stimuli were able to activate both pathways, whereas the K^+^-independent stimulus imiquimod strictly relied on the cage/MTOC-dependent pathway. Collectively, our results delineate two parallel and biologically distinct NLRP3 activation pathways, with smaller NLRP3 species employing an TGN/MTOC-independent NLRP3 activation pathway and decameric NLRP3 cages following the canonical TGN/MTOC-dependent NLRP3 activation pathway. Our results connect previous and at times conflicting results into a unified mechanistic framework of NLRP3 inflammasome activation. We hypothesize that these distinct pathway requirements may be relevant for effectively targeting NLRP3 and play a role in physiological and pathological relevant conditions of NLRP3 activity.

## Results

### NLRP3 inflammasome components reside in both cytosol and membranes before activation

The inactive NLRP3 oligomer is proposed to reside in TGN membranes with smaller species such as monomers described to be present in the cytosol ^13, 24, 25^ but the physiological relevance of the latter has not been fully explored. In this inactive state, it is still elusive where the next partners of the NLRP3 pathway, such as ASC, reside. Therefore, we decided to explore the subcellular localization of NLRP3 inflammasome components in frequently used, human macrophage-like cells, THP-1. Extracts from PMA-differentiated and LPS-primed or unprimed THP-1 WT and *NLRP3* knockout (KO) reconstituted with NLRP3 tagged with the bright monomeric fluorophore mNeongreen (mNG) were fractionated into P5 (heavy membranes), P100 (light membranes) and S100 (cytosol and vesicles) fractions by differential centrifugation ^20, 26^. Despite the expected fractionation of several well-known subcellular markers, NLRP3, ASC, NEK7, pro-caspase-1 and pro-IL-1β were detected in all fractions, with ASC and pro-caspase-1 being more abundant in the S100 (**Fig. 1A**). This distribution was neither modified between cell lines nor in the presence of LPS-priming (**Fig. 1A**). Our data suggested that critical NLRP3 inflammasome components reside in the same subcellular fractions, organelle-dependently and independently associated, highlighting the possibility that functional inflammasomes might arise at both locations.

**Figure 1.**
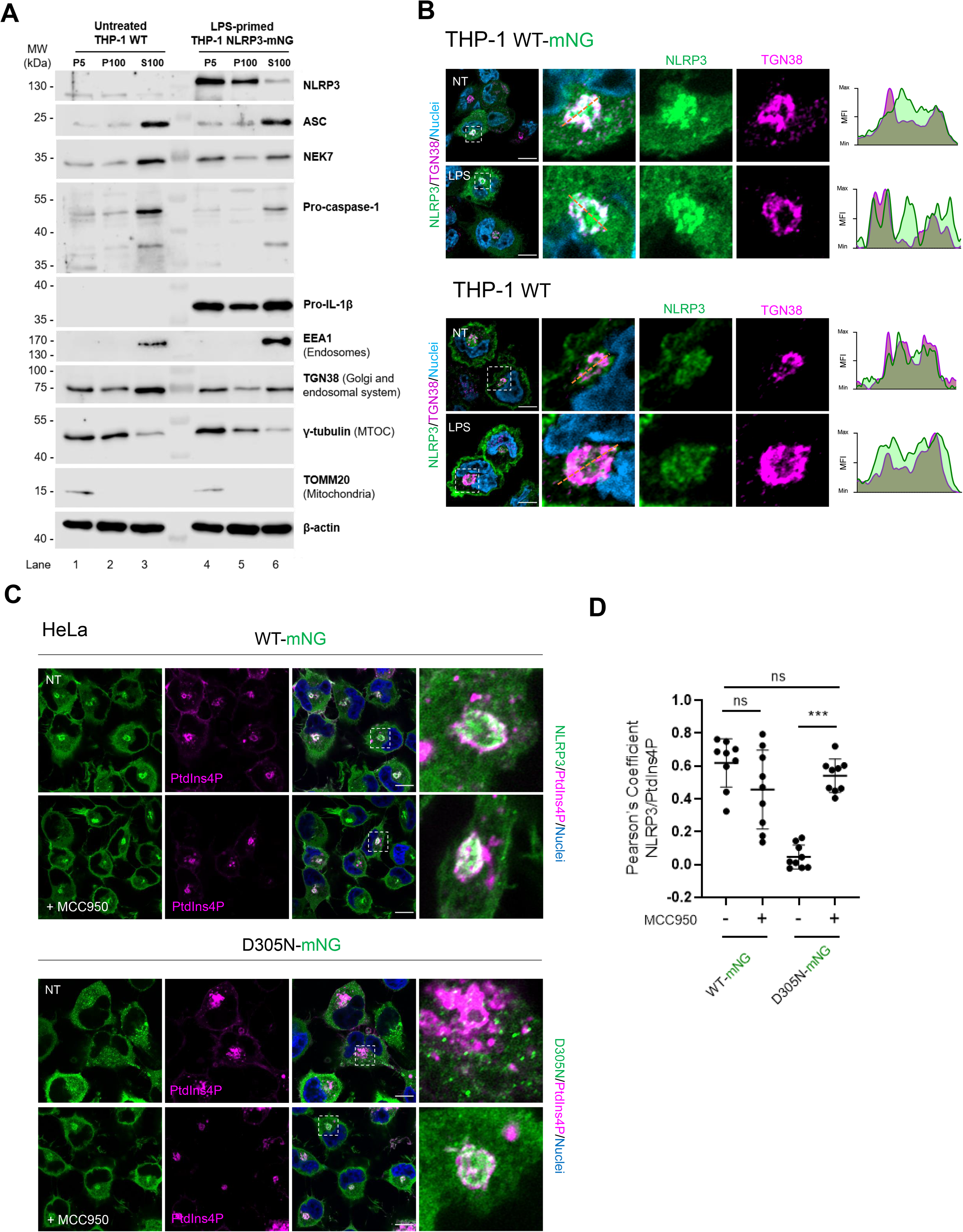
Inflammasome components reside in the cytosol and are associated with different organelles. **A)** Immunoblotting of extracts from untreated THP-1 WT or LPS-primed THP-1 *NLRP3* KO reconstituted with NLRP3-mNG cells for the indicated NLRP3 inflammasome components and organelle markers in P5 (heavy membrane), P100 (light membrane) and S100 (cytosol and vesicles) fractions. N=2 independent experiments. **B)** Representative immunofluorescence micrographs of THP-1 WT (upper) and NLRP3 *KO* cells reconstituted with NLRP3-mNG (lower panels) and primed or not (non-treated, NT) with LPS at 10 ng/ml for 4 h. Endogenous NLRP3 from THP-1 WT was stained with the specific NLRP3 antibody Cryo-2 (green). The TGN for both cell lines was stained with a specific TGN38 antibody (magenta) and nuclei with Hoechst 33342 (blue). Scale bar 10 µm, hashed boxes denote close-up regions. Representative line graphs depict changes in NLRP3 and TGN normalized mean fluorescence intensity (MFI). N=3-4 independent experiments. **C)** Representative micrographs of HeLa cells stably expressing WT NLRP3-mNG or CAPS mutant NLRP3 p.D305N-mNG transfected with the specific PtdIns4P sensor 2xSidM-mCherry (magenta) treated or not with MCC950 for 24 h. Nuclei were stained using Hoechst 33342 (blue). Scale bar 10 µm, hashed boxes denote close-up regions, N=3 independent experiments. **D)** Co-localization analysis of NLRP3-mNG or NLRP3 p.D305N-mNG and PtdIns4P in cells treated as in **C**. Values are represented by Pearsońs correlation coefficient with Costes automatic thresholding. Data are represented as mean ± SD and each data point represents one cell from three independent experiments (n=3 cells per independent experiment). ***p≤ 0.001, ns=not significant according to one-way ANOVA.

Microscopy experiments showed both NLRP3-mNG and endogenous NLRP3 unprimed or primed (with LPS) across the entire cell and strongly enriched in the TGN area before activation (**Fig. 1B)**, confirmed by line plot analysis (**Fig. 1B**). Previous results obtained by Andreeva *et al.,* ^13^ are in line with this finding, thereby excluding the possibility of the fluorophore preventing the formation of the oligomeric TGN-associated NLRP3. In addition, HeLa cells stably expressing NLRP3 WT-mNG were transfected with the phosphatidylinositol 4-phosphate (PtdIns4P) sensor-mCherry (2xSidM) that selectively labels PtdIns4P-rich membranes such as TGN, plasma membrane and endosomes ^27^. This sensor revealed NLRP3 residing with PtdIns4P (**Fig. 1C**), but also a pool of non-PtdIns4P-associated NLRP3. The NLRP3-PtdIns4P association was not affected by the presence of the specific NLRP3 inhibitor MCC950 (**Fig. 1C**), as confirmed by Pearson’s correlation analysis (**Fig. 1D**). Interestingly, the auto-active and CAPS disease-causing mutant NLRP3 D305N, when stably expressed in HeLa cells, spontaneously aggregated in small puncta in the cytosol that did not reside with the marker of PtdIns4P, coincident with a drastically lower correlation coefficient (**Fig. 1C, D**). However, after 24 hours (h) of treatment with MCC950, NLRP3 D305N was found at the TGN and correlated with PtdIns4P (**Fig. 1C, D**). These results led us to hypothesize that MCC950 can arrest NLRP3 in TGN membranes by stabilizing its cage structure, consistent with structural studies^14^. Moreover, the observation that key inflammasome components were present in non-TGN locations and that CAPS mutant NLRP3 formed oligomers distal to PtdIns4P and the TGN contributed to the notion that an activation pathway distal to TGN might exist.

### Non-decameric NLRP3 is not recruited to TGN membranes before activation

NLRP3 contains two adjacent polybasic regions described to be critical for membrane association: the KMKK motif (residues 131-134, e.g. blue box in **Fig. 2A**) and the polybasic region (residues 135-147, yellow box in **Fig. 2A**). The first motif has been reported to be dispensable for activation in human NLRP3 ^10^ but critical for mouse Nlrp3 ^20^. When taking a closer look at the decameric structure of human NLRP3, we observed that five of the second polybasic motifs are exposed on the same interface (**Fig. 2B**, dark blue alpha helices), which could be necessary for TGN binding, similar to how septin proteins for instance bind Golgi membranes using a clustering of polybasic motifs ^28^. We therefore hypothesized that disrupting the decamer formation would simultaneously prevent this concerted orientation of the polybasic regions and hence TGN binding. Interestingly, deletion of the N-terminal part of NLRP3 (PYD-linker domain) has been reported to lead to the formation of a hexamer instead ^24^, where only three polybasic regions are oriented more distally (**Fig. 2B**). In murine Nlrp3 such a deletion prevents cage formation and TGN recruitment ^13^. Although the membrane binding capabilities of the hexamer resulting from PYD+linker deletion were not investigated in human NLRP3, the deviation from the decamer orientation suggests that PYD and/or linker are critical for the organization of the decamer and its membrane association. Since the PYD domain is essential for signaling, we decided to solely delete the linker region (residues 95-134) in order to prevent decamer formation. This linker region is composed of 40 amino acids including the KMKK motif and is fully encoded by a single short exon in *NLRP3*, exon 3 (**Fig. 2A**). The NLRP3 structure appears to be modular and dictated by exons ^29^. We therefore anticipated exon 3 deletion to be tolerated by the protein. Especially the second polybasic region within the FISNA domain was left intact to fulfil its structural association to the nucleotide binding domain (NBD) and its critical role in changing the conformation of NLRP3 to allow for its activation ^25^ (**Fig. 2A, B**).

**Figure 2.**
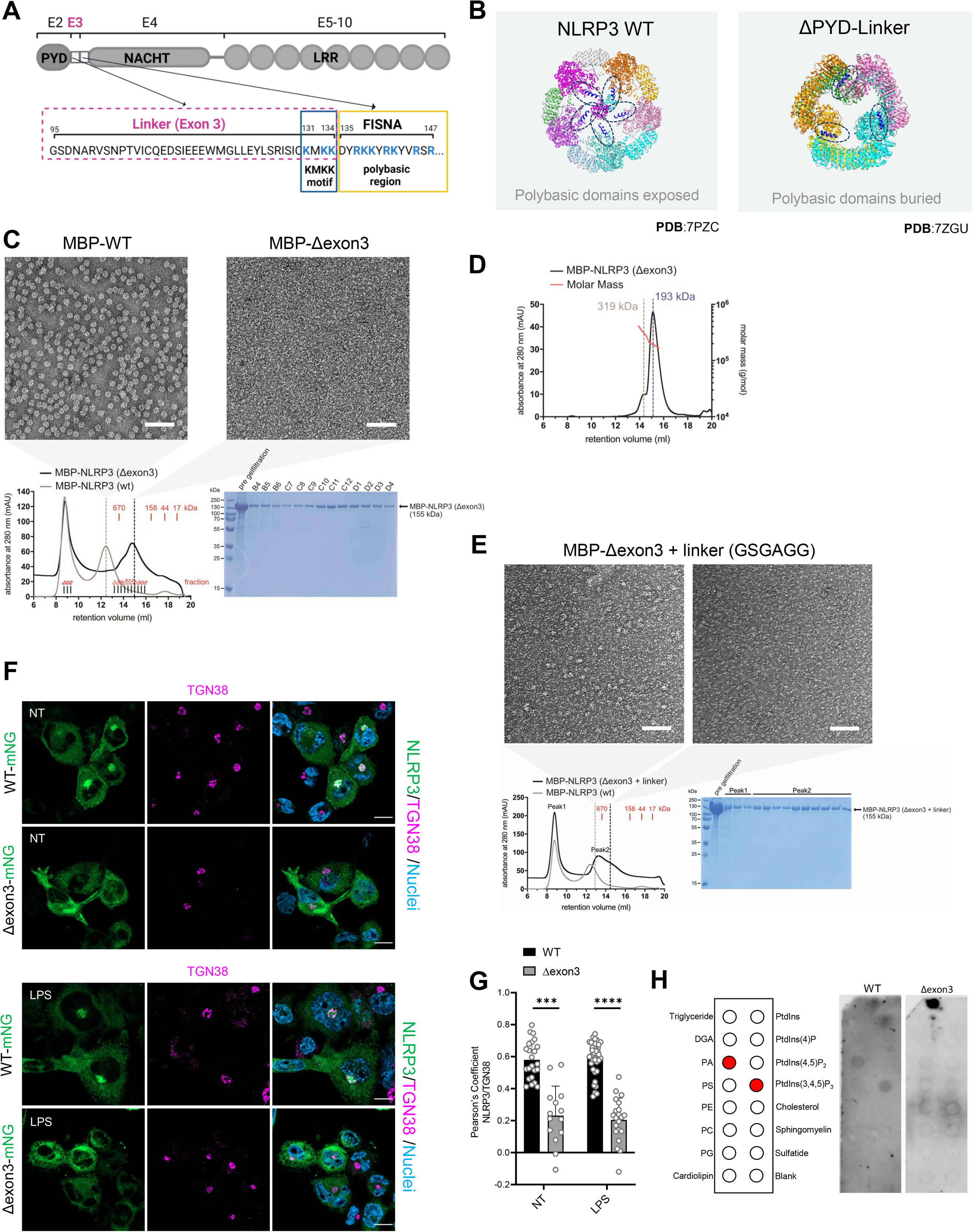
Δexon3 NLRP3 is organized in monomers unable to bind TGN membranes. **A)** Scheme of exons (E) and domain distribution of human NLRP3. Created with Biorender.com. **B)** Structure of the human double-ring NLRP3 decamer (left, pdb 7PZC) and human NLRP3 ΔPYD-linker hexamer (right, pdb 7VTP). Polybasic regions are highlighted and colored in dark blue. Different colors of the NLRP3 structures indicate different protomers. **C)** Negative-stain EM micrographs of the MBP-NLRP3 WT peak 2 (corresponding to the decamer) and the MBP-NLRP3 Δexon3 peak 2 (showing a lawn of NLRP3 particles smaller than the decamer). The scale bar for both is 100 nm. Below, SEC of recombinant human MBP-NLRP3 WT (grey trace) and MBP-NLRP3 Δexon3 (black trace) is shown. Expected molecular weights based on a gel filtration standard and elution fractions of purified MBP-NLRP3 Δexon3 are indicated in red. The indicated fractions were analyzed by SDS-PAGE (Coomassie staining). **D)** SEC-MALS of recombinant human MBP-NLRP3 Δexon3 peak 2 on a Superose 6 Increase 10/300 GL column. The estimated molar masses are indicated by the red line. The calculated mass of the MBP-NLRP3 Δexon3 monomer is 154.8 kDa. **E)** Negative-stain EM micrographs of the left and right part of the peak 2 of MBP-Δexon3 NLRP3 + GSGAGG linker. The left part of peak 2 already shows a more pronounced capability to form higher-order oligomers in the presence of GSGAGG linker. The scale bar for both is 100 nm. SEC as in **C** of MBP-NLRP3 WT (grey peaks) and MBP-Δexon3 NLRP3 + GSGAGG linker (black peaks). On the right, Coomassie-stained SDS-PAGE analysis of the SEC run of MBP-Δexon3 NLRP3 + GSGAGG linker. **F)** Representative immunofluorescence micrographs of LPS primed or not (NT) THP-1 *NLRP3* KO cell line reconstituted with NLRP3 WT- or NLRP3 Δexon3-mNG (both in green).TGN stained with a specific TGN38 antibody (magenta) and nuclei with Hoechst 33342 (blue). Scale bar 10 µm. N=4 independent experiments. **G)** Co-localization levels between WT or Δexon3 NLRP3 and TGN38 in cells treated as in **F**. Values are represented by Pearsońs correlation coefficient with Costes automatic thresholding. Each bar represents the mean ± SD and each datapoint represent single cells combined from 3 independent experiments. ***p≤ 0.001; ****p≤ 0.0001 according to one-way ANOVA. **H)** *In vitro* lipid strip assay of the purified human NLRP3 WT decamer and NLRP3 Δexon3 monomer. On the left, the arrangement of different lipids on the membrane is shown, with lipids bound by NLRP3 WT highlighted in red. N= 3 for WT and N=2 for Δexon3 independent experiments.

To explore whether NLRP3, when lacking the exon 3 linker region (termed Δexon3), was able to assemble into decamers, human NLRP3 Δexon3 protein was expressed in baculovirus-infected *Sf9* insect cells as a fusion protein with maltose-binding protein (MBP), and purified to homogeneity as before ^14^. Size-exclusion chromatography (SEC) of this protein revealed two elution peaks: one close to the void volume, indicative of an undefined high-molecular-weight aggregate (previously observed also for the NLRP3 WT protein ^14^), and a wider one covering a size from 158 to 670 kilodaltons (kDa) (**Fig. 2C**) indicating that Δexon3 NLRP3 could form monomers to tetramers, but not hexamers as we expected ^24^. The negative-stain electron microscopy (EM) images of MBP-Δexon3 NLRP3 did not show a clear and defined structure when compared to WT, but clearly confirmed smaller assemblies and the absence of the typical decamer observed for WT protein (**Fig. 2C**). Consistently, SEC coupled with multi-angle light scattering (SEC–MALS) experiments revealed an average molecular mass in the range of 193-319 kDa for peak 2, which corresponds to monomers or dimers of the MBP– Δexon3 NLRP3 protein (**Fig. 2D)**. These results indicated that Δexon3 NLRP3 cannot be organized in the typical NLRP3 decameric structure suggesting that the spatial and conformational flexibility provided by this linker region is needed to conform the decamer.

To elucidate whether the specific linker sequence encoded by the exon 3 is directly needed for the formation of the decamer or whether it functions more as a coordinator/facilitator of the assembly we substituted the 40 amino acids of the linker sequence with the short flexible linker sequence (GSGAGG). The sole presence of these six amino acids enabled NLRP3 to form slightly bigger, albeit still non-decameric, species (**Fig. 2E**). As the Δexon3 NLRP3 protein appeared soluble and fully folded when purified, we were intrigued to explore the subcellular localization of the Δexon3 non-cage form of NLRP3 in cells. Microscopy studies on PMA-differentiated and LPS-primed or unprimed THP-1 *NLRP3* KO cells reconstituted with NLRP3 WT- or Δexon3-mNG, showed that in the absence of the linker, NLRP3 decreased its localization at the TGN as expected (**Fig. 2F**, quantified in **G**). Similarly, in HeLa cells stably expressing Δexon3 NLRP3, a lower colocalization with the TGN membranes was apparent in resting cells as visualized via the TGN46 marker (**Fig. S1A, B** non-treated (NT) conditions). The binding to the TGN is related to the last four amino acids of the NLRP3 linker motif encoded by the exon 3 ^20^ (KKKK in mice and KMKK in humans, blue box in Fig. 2A). However, in resting cells, removal of positive charges in a human NLRP3 KMKK>AMAA mutated construct (NLRP3 AMAA) was still able to localize to the TGN (**Fig. S1A, B** non-treated (NT) conditions). To further show that the KMKK motif was dispensable for human NLRP3 activation, HeLa cells stably expressing NLRP3 AMAA or Δexon3 NLRP3, both oligomerized and formed specks similar to NLRP3 WT upon nigericin stimulation (**Fig. S1A** nigericin pictures). However, HeLa cells expressing NLRP3 WT or NLRP3 AMAA presented a decreased colocalization with TGN after activation compared to unstimulated cells, but cells expressing Δexon3 NLRP3 protein showed no colocalization between NLRP3 and TGN (**Fig. S1A, B** nigericin conditions). Our data suggest that, unlike the KMKK motif, the second polybasic region is sufficient for human NLRP3 to localize to membranes. However, since the Δexon3 construct still preserves the second polybasic region, we conclude that this polybasic region only allows for membrane lipid association in a decameric arrangement.

This was, indeed, confirmed by lipid strip binding assays of the purified human Δexon3 NLRP3 protein. Whereas human WT NLRP3 bound phosphatidic acid (PA) and PtdIns-(3,4,5)-P_3_, Δexon3 NLRP3 protein lost the ability to bind these and any other lipids (**Fig. 2H**). These results further suggest that the NLRP3 linker sequence encoded by the exon 3 may provide the proper spatial arrangement necessary for the polybasic motif exposure and TGN recruitment observed in resting conditions.

Curiously, purified even the human NLRP3 WT protein was unable to bind other phosphoinositides such as PtdIns4P previously described for mouse Nlrp3 ^13, 24^, potentially highlighting the differences in membrane affinity between both species. To further explore this difference of PtdIns4P binding affinity, HeLa cells stably expressing NLRP3 WT were transfected with the PtdIns4P marker 2xSidM and treated with nigericin. We observed a decrease in colocalization of PtdIns4P with the active NLRP3 oligomers **(Fig. S1C, D**) as we showed above for the autoactive D305N NLRP3 mutation (**Fig. 1D, E**). This loss in colocalization was further maintained even when the cage conformation was enforced by MCC950 addition **(Fig. S1C, D**). In conclusion, the human NLRP3 decamer is recruited to TGN not necessarily through PtdIns4P association but rather through binding other lipids. In contrast to WT NLRP3, the Δexon3 NLRP3 cannot not be organized in a lipid binding decamer.

### Non-decameric NLRP3 forms TGN- and MTOC-independent specks in response to nigericin

Our previous data indicated that Δexon3 NLRP3 might provide a unique opportunity to study the functionality of an NLRP3 inflammasome that is cage- and membrane binding-independent. We therefore first assessed whether the Δexon3 NLRP3 could still be activated in macrophage cells. LPS-primed THP-1 cells stably expressing NLRP3 WT- or Δexon3-mNG treated with nigericin confirmed both cell lines to be able to form NLRP3 specks, a hallmark of functional inflammasomes (**Fig. 3A**). Interestingly, we realized that while Δexon3 NLRP3 only formed TGN-independent specks, WT NLRP3 presented both, TGN-dependent and independent specks formation (**Fig. 3A**, quantified in **B**), sometimes in the same cell. That active inflammasomes may originate from TGN-localized cages has been shown before ^13^, but the observation of TGN-membrane-independent NLRP3 specks posed the intriguing question, whether these were also competent inflammasomes.

**Figure 3.**
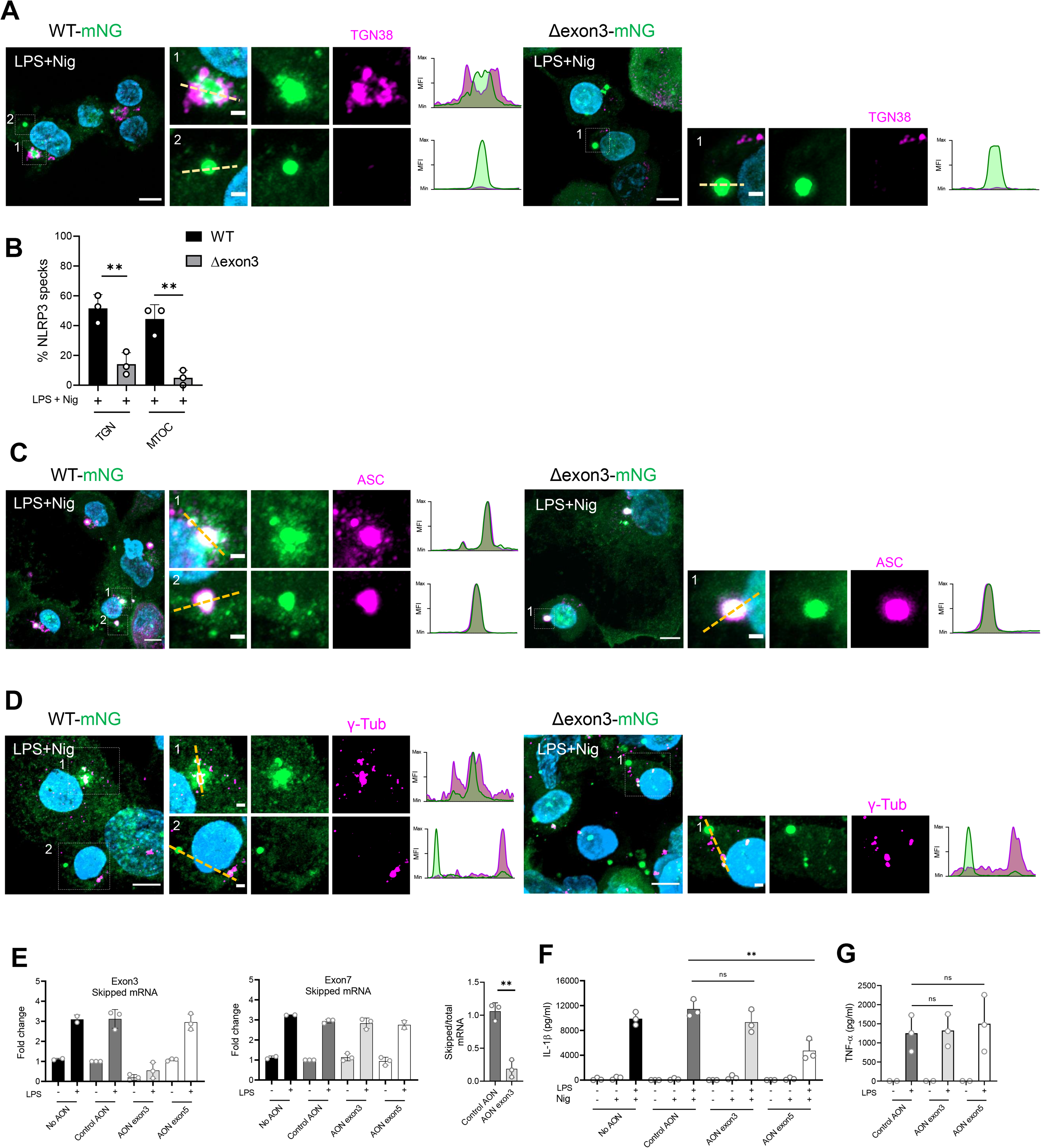
Δexon3 NLRP3 forms a functional inflammasome and constitutes a tool to study membrane-independent NLRP3 activation. **A)** Representative immunofluorescence micrographs of the indicated LPS-primed and nigericin-stimulated THP-1 cell lines. TGN was stained with the TGN38 antibody (magenta) and nuclei with Hoechst 33342 (blue). Scale bar 5 µm, 1 µm for close-ups. Representative line graphs depicting changes in NLRP3 and TGN normalized mean fluorescence intensity (MFI) are shown. N=3 independent experiments **B)** Percentage (%) of membrane (TGN) and (MTOC)-associated NLRP3 specks (WT or Δexon3) of the microscopy data examples shown in **A** and **D**. Each bar represents the mean ± SD and each datapoint represents the average of 3 tiles (3×3 FOV) (200 cells approx.). N=3 independent experiments. **p≤ 0.01 according to one-way ANOVA. **C)** As in **A** but endogenous ASC marked with a specific ASC antibody (magenta). NLRP3 and ASC representative line graphs depicting changes in normalized mean fluorescence intensity (MFI) are shown. N=3 independent experiments. **D)** As in **A** but MTOC marked with the γ-tubulin antibody (magenta). NLRP3 and MTOC representative line graphs depicting changes in normalized mean fluorescence intensity (MFI) are shown. N=3 independent experiments. **E)** RT-qPCR showing endogenous skipping of the *NLRP3* exon 3 (linker region) in PMA-differentiated THP-1 WT cells. Upon exon skipping for 24 h, the cells were primed or not with LPS. Left, fold change of *NLRP3* exon 3 mRNA expression. Middle, fold change of *NLRP3* exon 7 mRNA expression used as a negative control. Right, ratio of skipped exon 3 compared to total NLRP3 mRNA (using exon 7 as control). **p≤ 0.01 according to Student’s t-test. N=3 independent experiments. **F)** IL-1β release (triplicate ELISA, mean + SD) from THP-1 WT treated as in **E** and further stimulated or not with nigericin. Skipping of the *NLRP3* exon 5 was used as a positive control of dampened NLRP3 activation. **p≤ 0.01; ns, non-significant according to one-way ANOVA. N=3 independent experiments (combined). **G)** TNF-α release (triplicate ELISA, mean + SD) from THP-1 WT treated as in **F**. N=3 independent experiments (combined).

To form an active NLRP3 inflammasome, NLRP3 must engage ASC, the next partner in the pathway, which culminates in NLRP3-ASC speck formation ^1^. As expected, upon nigericin stimulation, LPS-primed THP-1 cells stably expressing WT or Δexon3 NLRP3 both formed NLRP3-ASC specks (**Fig. 3C**). In THP-1 WT NLRP3 cells, we again observed two clear phenotypes for NLRP3-ASC distribution: NLRP3-ASC platforms surrounded by multiple smaller NLRP3-ASC puncta, resembling the TGN staining imaged above (**Fig. 3A, C** NLRP3 WT example 1); and single, well-delineated NLRP3-ASC platforms distal to TGN staining (**Fig. 3A, C** NLRP3 WT example 2). However, in the case of THP-1 Δexon3 NLRP3, only the latter type of NLRP3-ASC platform, the TGN distal one, could be observed (**Fig. 3A, C**). When comparing the single TGN-distal NLRP3-ASC platforms from WT vs Δexon3 NLRP3-expressing cells, no differences were detected between them in terms of fluorescence intensity and area (**Fig. S2A**). Interestingly, we also confirmed that ASC organized in a platform surrounded with multiple and smaller ASC puncta was also surrounded by TGN membranes, whereas the ASC single well-delineated platform was typically not (**Fig. S2B**). Importantly, we also located ASC speck formation surrounded by Golgi membranes in human primary macrophages upon LPS and nigericin treatment (**Fig. S2C**), showing they occur at endogenous levels even in primary cells. In addition, nigericin activation yielded similar caspase-1 cleavage levels for the WT and Δexon3 NLRP3; further proving the functionality of the Δexon3 NLRP3 inflammasome in reconstituted THP-1 cells (**Fig. S3A, B**).

As the MTOC was proposed as the final destination of NLRP3 to culminate in the formation of an active inflammasome ^23^, we also performed immunofluorescence analysis of the MTOC in LPS-primed THP-1 cells reconstituted with WT or Δexon3 NLRP3 by using the specific MTOC marker, γ-tubulin ^30^. Again, nigericin stimulation produced two clear phenotypes for NLRP3 WT specks: MTOC-dependent and MTOC-independent NLRP3 specks (**Fig. 3D**, quantified in **B)**. However, as expected and following the line of our previous results, Δexon3 NLRP3 specks were all MTOC-independent (**Fig. 3D**). Thus, WT NLRP3 seemed to be able to form both TGN/MTOC-dependent and -independent NLRP3 specks upon nigericin activation, whereas Δexon3 NLRP3 only formed TGN/MTOC-independent ones. This led us to hypothesize that the NLRP3 decamer by its spatial arrangement of polybasic motifs should be associated with the described TGN/MTOC-dependent inflammasome activation pathway ^13, 24, 25^, whereas the smaller NLRP3 species might activate and form specks via a separate, membrane-independent pathway in the cytosol as a signalling competent environment (*cf.* Fig. 1A).

To prove that the observation of two types of functional NLRP3 pathways was not a consequence of an NLRP3 overexpression phenomenon, we endogenously disrupted the expression of the NLRP3 linker sequence by skipping the *NLRP3* exon 3 in THP-1 cells using an intra-exonic anti-oligonucleotide (AON), similar to an approach recently used to skip the exon 5 ^29^. The effects of the AON to target exon 3, a negative control AON and an AON targeting exon 5 were tested in human THP-1 cells by qPCR. Exon 3 was specifically targeted and skipped by about 80% without affecting other NLRP3 exons like exon 7 and hence total mRNA NLRP3 levels (**Fig. 3E**). Strikingly, in terms of IL-1β release no significant differences were detected whether NLRP3 exon 3 was skipped or not, while exon 5 skipping dampened IL-1β production as expected ^29^ (**Fig. 3F**). In addition, TNF-α release was not affected by any exon-skipping approaches (**Fig. 3G**). These results indicated that an NLRP3 protein incapable of organising in a decamer structure was still able to form a functional inflammasome measured by ASC assembly, caspase-1 cleavage and IL-1β production. This further confirmed that both decameric and non-decameric NLRP3 species can form active NLRP3 inflammasomes.

### Microtubule-dependent NLRP3 specks accelerate inflammasome activation

Having confirmed that Δexon3 NLRP3 is functional in cells, we went on to study in detail the microtubule dependence of NLRP3 by comparisons with WT NLRP3. We performed highly resolved live cell imaging experiments in LPS primed and nigericin stimulated THP-1 cells stably expressing WT- or Δexon3-mNG NLRP3 using the highly specific probe for microtubules, SIR-tubulin ^31^. Here, only the WT NLRP3 speck was able to consistently associate with the MTOC, while the Δexon3 NLRP3 speck appeared in different cellular locations, presumably free in the cytosol, as the signal was completely MTOC-independent (**Fig. 4A**, **S4A** WT NLRP3 speck in the MTOC example 2 and **supplementary movies 1 and 2**). Indeed, we could also visualize the formation of an MTOC-independent WT NLRP3 speck, indicating the dynamic presence of both types of specks, and hence pathways, in the case of WT NLRP3 (**Fig. S4A** WT NLRP3 MTOC-independent speck and **supplementary movie 3**). Taking a closer look at the WT NLRP3, we observed NLRP3 to be recruited to endosomal-like structures upon nigericin stimulation which trafficked and collapsed at the MTOC by 25 min, resulting in NLRP3 speck formation (**Fig. 4A**). We noted that, the MTOC-dependent WT NLRP3 specks formed faster (**Fig. 4A, B**), while MTOC-independent specks formed more slowly, i.e. within 55 min. Interestingly, there were no differences in the time to speck formation for the WT and Δexon3 NLRP3 MTOC-independent specks (**Fig. 4B**). This phenomenon observed for WT NLRP3 also correlated with an accelerated cell death measured by real-time SYTOX orange uptake imaging (**Fig. 4C**), faster ASC speck formation (**Fig. 4D**) and earlier IL-1β release (**Fig 4E**), that was abolished in the presence of MCC950 (**Fig. 4D, E**). No differences were detected in TNF-α release between THP-1 *NLRP3 KO* or reconstituted with Δexon3 NLRP3, but we always detected a higher TNF-α release in *KO* cells reconstituted with WT NLRP3 (**Fig. 4F**). Additionally, we explored the mobility of both MTOC-dependent and - independent NLRP3 specks by single object tracking analysis. We observed that the MTOC-independent specks clearly showed increased mobility (**Fig. S4B)**, consistent with their independence of membranes and microtubules. If this was the case, MTOC-independent NLRP3 speck formation should not be sensitive to microtubule depolymerizing agents ^23^.

**Figure 4.**
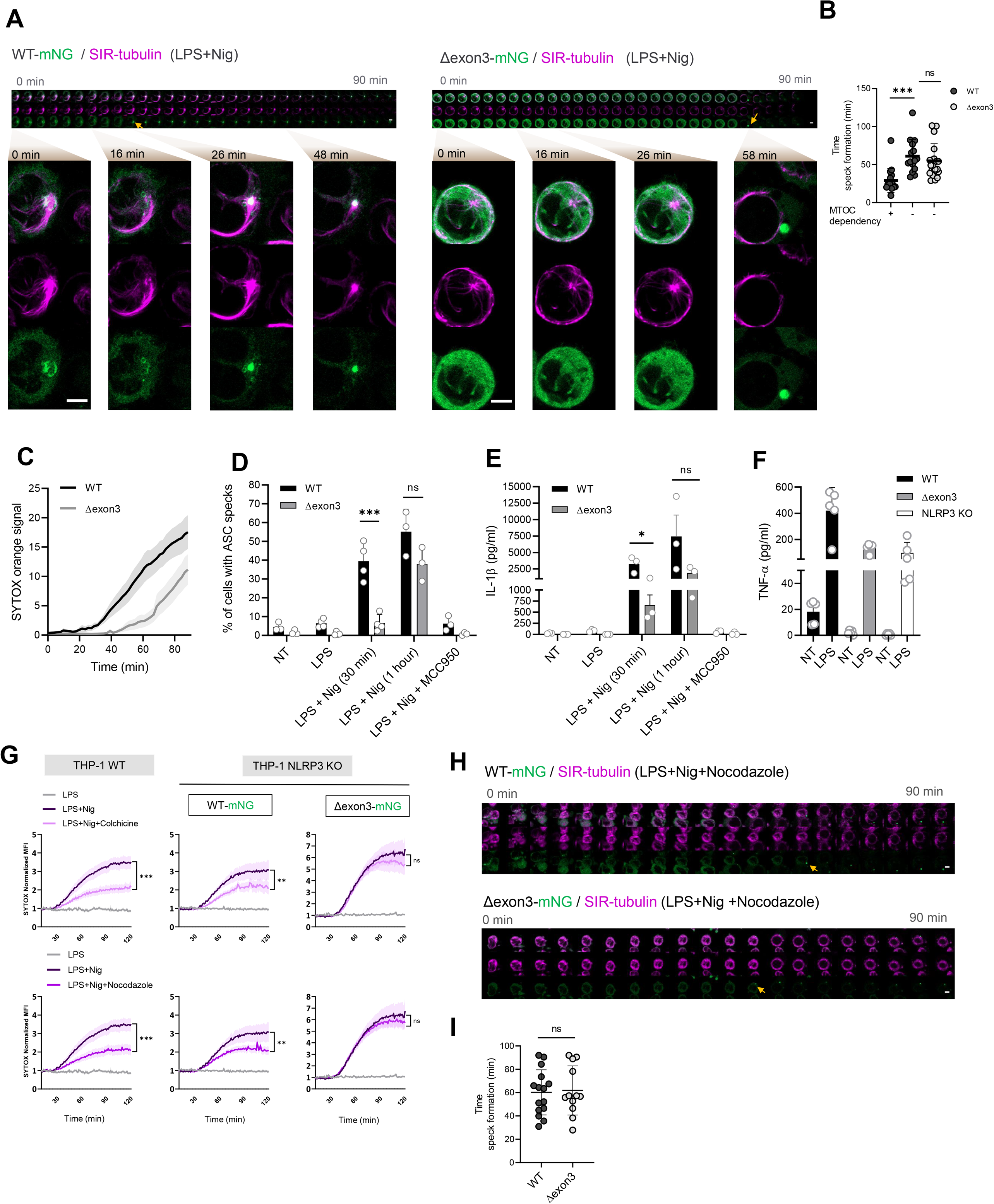
Membrane-associated NLRP3 forms the speck in the MTOC accelerating inflammasome activation. **A)** Representative live cell imaging timeline of the indicated LPS-primed and nigericin-stimulated THP-1 cell lines. Microtubules were stained with the specific probe SIR-tubulin (magenta). Shown are 30 frames from a time-lapse sequence, spaced 2.4 min apart. Arrows denote first occurrence of fully formed NLRP3 speck. Close-up images at 0, 16, 26 and 48 or 58 min after nigericin addition are shown. Scale bar 5 µm. N=3 independent experiments. **B)** Time of MTOC-dependent and -independent speck formation quantified in cells treated as in **A**. Each bar represents the mean ± SD and each datapoint represents one speck. ***p≤ 0.001; ns, non-significant according to Kruskal-Wallis test with Dunn’s multiple comparisons test. N=3 independent experiments. **C)** Cell death in time measured by SYTOX orange uptake treated as in **A.** N=3 independent experiments. **D)** Percentage (%) of ASC speck formation from the indicated THP-1 cell lines upon the treatments shown. Each bar represents the mean ± SD and each datapoint represents the average of 3 tiles (3×3 FOV) (200 cells approx.). ***p≤0.001; ns=non-significant, according to one-way ANOVA. N=3-4 independent experiments. **E)** IL-1β (triplicate ELISA, mean + SD) release as treated in **D.** *p≤0.05; ns=non-significant, according to Student’s t-test. N=3 independent experiments (combined). **F)** TNF-α release (triplicate ELISA, mean + SD) from the indicated THP-1 cell lines LPS-primed or not. N=5 independent experiments (combined). **G)** Kinetics of cell death from the indicated THP-1 cell lines upon the treatments shown. SYTOX orange uptake is represented as the mean fluorescence intensity (MFI) ± SEM normalized to the signal of the first 5 min of each condition. N=3 independent experiments. ***p≤ 0.001; **p≤ 0.01; ns, non-significant according to one-way ANOVA. **H)** Live cell imaging as in **A** but including the microtubule disrupting agent nocodazole. Shown are 20 frames from a time-lapse sequence, spaced 4.6 min apart. Arrows denote first occurrence of fully formed NLRP3 speck. Scale bar 5 µm. N=3 independent experiments. **I)** Quantification of MTOC-dependent and independent specks. Each dot represents one speck from 2 independent experiments. ns=non-significant, according to Student’s t-test.

The introduction of two general microtubule disruptors, colchicine or nocodazole, indeed did not affect the cell death associated with Δexon3 NLRP3 inflammasome activation, while WT NLRP3 expressing cells were clearly and significantly affected (**Fig. 4G**). Moreover, similar results were obtained upon trafficking disruption via specific inhibition of dynein (using ciliobrevin D) or HDAC6 (using ricolinostat) (**Fig. S4C**). Live cell imaging experiments in the presence of nocodazole clearly showed the formation of the MTOC-independent WT NLRP3 speck in a completely disrupted microtubule system (**Fig. 4H** and **supplementary movie 4**) that occurred at a similar time compared to the Δexon3 NLRP3 speck formation (**Fig. 4H** quantified in **4I**). This indicates that only the MTOC-associated NLRP3 specks are responsible of the accelerated inflammasome activation. Interestingly, microtubule inhibition before nigericin treatment led to a WT NLRP3 trapped in non-travelling endosomal-like vesicles over 20 min (**Fig. S4D**). These results demonstrate the existence of two parallel and biologically distinct NLRP3 activation pathways: a previously described MTOC-dependent one, directly related to the NLRP3 decamer; and a second MTOC-independent one, related to non-decameric NLRP3 species. Additionally, we show that, although NLRP3 cytosolic specks are entirely functional upon nigericin stimulation, the MTOC-associated ones form earlier and execute inflammasome functions like IL-1β release and cell death faster. Thus, the existence of the decamer and/or the association with the structural framework of the MTOC-dependency may act as a catalyst for rapid assembly of NLRP3 platforms that results in a faster inflammasome activation response.

### Potassium efflux is critical for the microtubule-independent, imiquimod for the microtubule-dependent NLRP3 activation pathway

A question arising from these results was, whether different stimuli might favor one pathway over the other. Therefore, using LPS-primed THP-1 cells stably expressing WT or Δexon3 NLRP3-mNG, we explored whether the MTOC-dependent and -independent NLRP3 activation pathway previously described for nigericin, could be also extrapolated to other potassium-dependent stimuli and to imiquimod, an NLRP3 activator considered to be potassium-independent. The introduction of other NLRP3 potassium-dependent activation stimuli such as the monosodium urate (MSU) crystals ^32^ and the *Staphylococcus aureus* leukocidin A/B (LukAB) ^33^ also induced IL-1β release and cell death in WT and Δexon3 NLRP3 that were efficiently blocked in the presence of high extracellular KCl (**Fig. 5A, B**). Thus, potassium-dependent stimuli were able to activate both MTOC-dependent and -independent NLRP3 pathways. Interestingly, THP-1 cells expressing Δexon3 NLRP3 treated with imiquimod presented a damped IL-1β release and cell death (**Fig.5A, B**) that were completely abolished in the presence of high extracellular KCl, indicating that, in agreement with recent results in human cells ^34^, the residual IL-1β in this system was still related to K^+^ efflux (**Fig 5A**). Besides, nigericin activation led to pro-caspase-1, GSDMD, and pro-IL-1β cleavage and their release to a similar extent for WT and the Δexon3 NLRP3 variant (**Fig. 5C**). Conversely, caspase-1 and IL-1β cleavage and release and ASC speck formation were strongly reduced for Δexon3 NLRP3 upon imiquimod stimulation (**Fig. 5C-E**). These results reveal the inability of an MTOC-independent NLRP3 to respond to imiquimod. This observation was in line with microscopy experiments showing that, unlike nigericin (*cf.* **Fig. 3**), WT NLRP3 formed mostly TGN-associated specks upon imiquimod stimulation (**Fig. 5F**) that were further MTOC-associated (**Fig. 5G**). Consistently, imiquimod-induced cell death was significantly more sensitive to the microtubule-depolymerizing drug nocodazole (**Fig. 5H**). These results indicate that imiquimod stimulation exclusively relies on the NLRP3 cage/TGN/MTOC-dependent pathway while the NLRP3 TGN/MTOC-independent pathway plays a role exclusively for K^+^-dependent stimuli.

**Figure 5.**
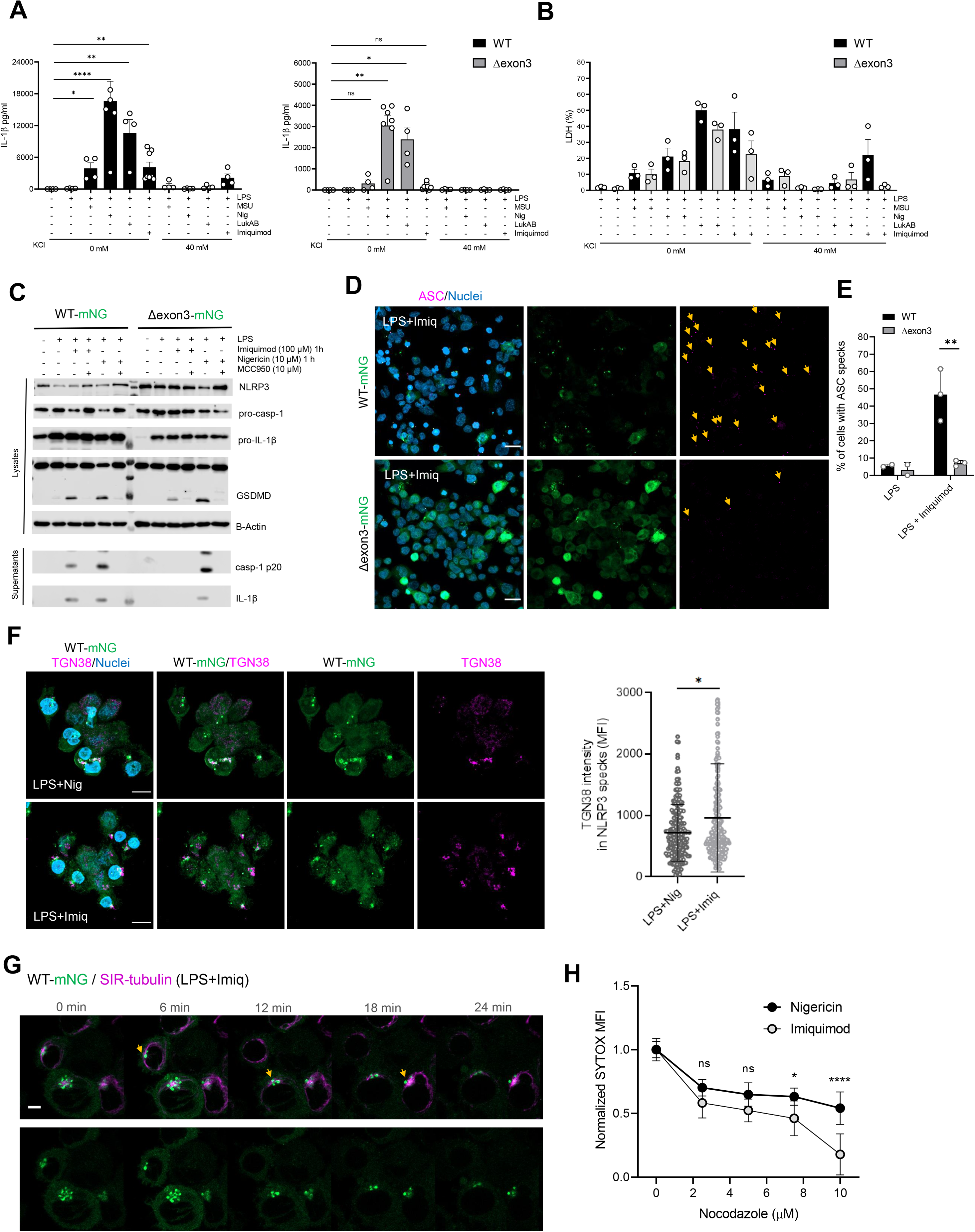
K^+^ efflux-activating stimuli are responsible for the MTOC-independent NLRP3 activation whereas imiquimod only induces the MTOC-dependent NLRP3 activation. **A, B)** IL-1β and LDH release (performed in triplicates, mean + SD) of the indicated THP-1 cell lines stimulated with different NLRP3 inflammasome inducers in low and high K^+^ conditions as indicated in the graphs. ****p≤ 0.0001; **p≤ 0.01; *p≤ 0.05; ns, non-significant according to one-way ANOVA. N=3-7 independent experiments (combined). **C)** One representative of three immunoblots from cells treated as indicated in the image. **D)** Representative confocal microscopy images of ASC speck formation in LPS-primed THP-1 cells in the presence or absence of imiquimod. Scale bar 20 µm. Arrows denote occurrence of fully formed ASC speck. N=3 independent experiments. **E)** Percentage (%) of ASC speck formation from **D**. *p≤0.05 according to Student’s t-test. N=3 independent experiments. **F)** Representative immunofluorescence micrographs of the indicated LPS-primed and nigericin- or imiquimod-stimulated THP-1 cell lines. TGN marked with the TGN38 antibody (magenta) and nuclei with Hoechst 33342 (blue). Scale bar 10 µm. Quantification of TGN38 MFI in NLRP3 specks. Each dot represents one speck from 3 independent experiments. *p≤ 0.05; according to Mann*–*Whitney U test. **G)** Representative live cell imaging of LPS-primed THP-1 NLRP3 WT-mNG upon imiquimod stimulation. Microtubules are marked with the specific probe SIR-tubulin (magenta). N=3 independent experiments. **H)** Cell death measured by SYTOX orange uptake in THP-1 cells treated with nigericin or imiquimod in the presence of different concentrations of nocodazole. ****p≤ 0.0001; *p≤ 0.05; ns, non-significant according to two-way ANOVA. N=3 independent experiments.

### Microtubule-dependent NLRP3 activation leads to TGN aggregation in an ASC dependent manner

Recently, cellular membranes have emerged as pivotal players in NLRP3 inflammasome functionality ^5, 12^. Especially TGN dispersion was proposed to play a major role in NLRP3 activation ^13, 20, 22, 23^. Having obtained evidence for two parallel pathways of which only one was associated with the TGN, we sought to explore to which extent TGN dispersion correlated with TGN/MTOC-dependent and TGN/MTOC-independent NLRP3 activation phenotypes. Interestingly, upon imiquimod stimulation we did not observe a clear TGN dispersion in WT NLRP3-mNG but rather an aggregation as indicated by TGN increased fluorescence intensity (**Fig. 6A, B**). Additionally, TGN aggregation was not observed in the case of Δexon3 NLRP3-mNG, indicating that this phenomenon is dependent on TGN-located NLRP3 (**Fig. 6A, B**). Upon nigericin stimulation, both phenotypes of cells – dispersion and aggregation (measured by TGN intensity increase and TGN area, respectively) – were observed for WT NLRP3, whereas only dispersion was detected for Δexon3 NLRP3 (**Fig. 6A, B**), indicating that in the absence of NLRP3 on the TGN only dispersion appears to occur. Furthermore, THP-1 cells with endogenous NLRP3 expression also exhibited TGN aggregation with imiquimod and both, dispersion or aggregation, with nigericin stimulation (**Fig. 6C**). MCC950 treatment did not abolish TGN dispersion but interestingly, strongly reduced TGN aggregation (**Fig. S5A**), indicating further that the latter is NLRP3-dependent.

**Figure 6.**
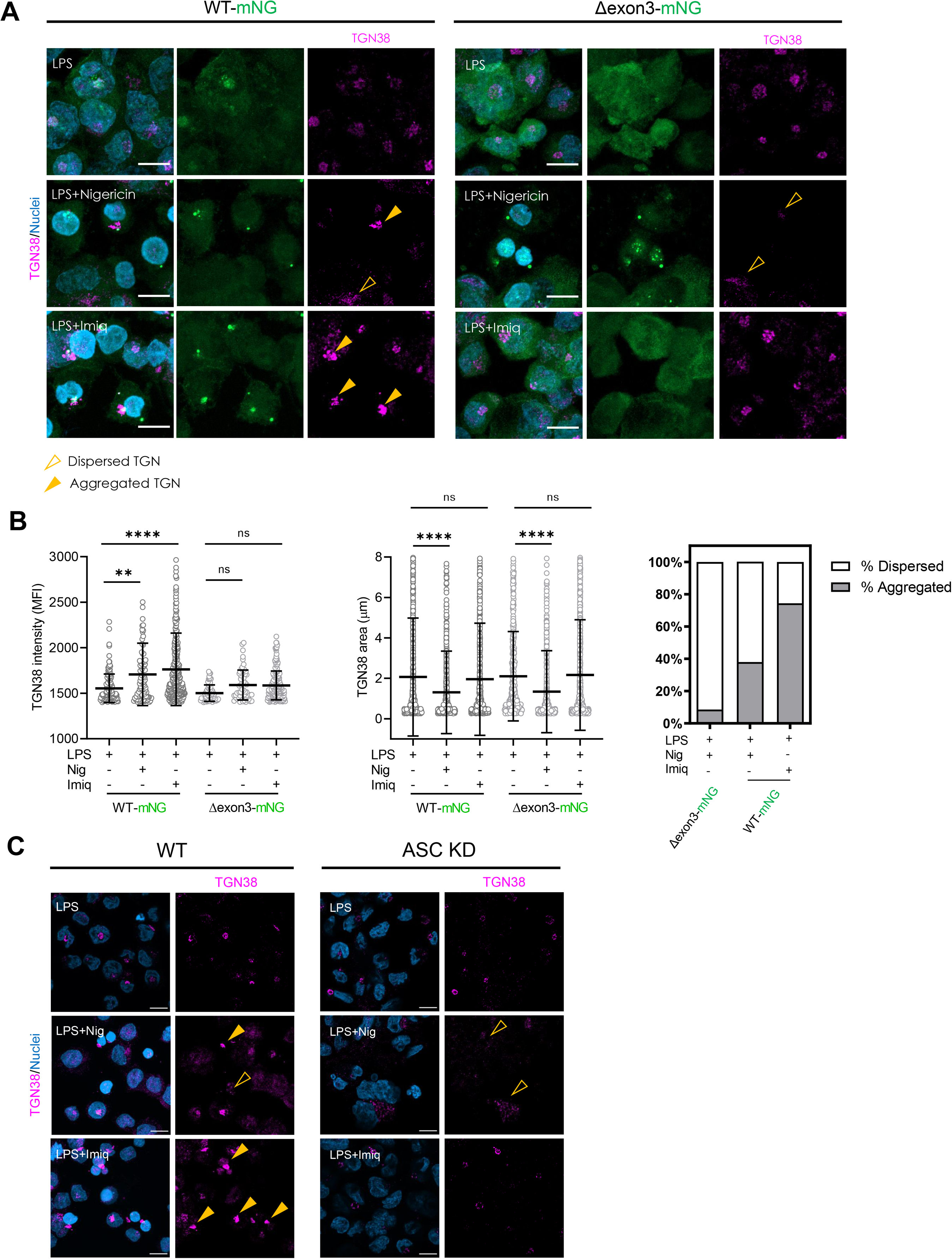
Membrane-bound NLRP3 leads to membrane aggregation in an ASC-dependent manner. **A)** Representative immunofluorescence micrographs of the indicated LPS-primed and nigericin- or imiquimod-stimulated THP-1 cell lines. TGN stained with the TGN38 antibody (magenta) and nuclei with Hoechst 33342 (blue). Filled triangles denote TGN aggregation, open triangles, TGN dispersion. Scale bar 10 µm. N=3 independent experiments. **B)** Quantification of the TGN aggregation (measured by the TGN mean fluorescence intensity, MFI, left), TGN dispersion (measured by the TGN area, middle) and percentage (%) of TGN dispersed vs aggregated (right) of **A**. ****p≤ 0.0001; **p≤ 0.01; *p≤ 0.05; ns, non-significant according to one-way ANOVA. N=2-3 independent experiments. **C)** Representative immunofluorescence micrographs of LPS-primed and nigericin- or imiquimod-stimulated THP-1 WT and *ASC* KD cell lines. TGN marked with the TGN38 antibody (magenta) and nuclei with Hoechst 33342 (blue). Scale bar 10 µm. N=3 independent experiments.

This first reported TGN aggregation in the context of inflammasome activation led us to consider whether other inflammasome components might also be involved. We noticed that most of the studies on TGN dispersion were performed either in non-macrophage cells not expressing ASC, or in an *ASC* KO background, or in the presence of an NLRP3 arrested by the inhibitor MCC950 ^20, 22, 23, 35^, excluding the study of the potential impact, if any, of ASC on the NLRP3-TGN interface. In line with the abovementioned published studies, we observed clear TGN dispersion in THP-1 ASC knockdown (KD) cells treated with LPS plus nigericin or imiquimod, (**Fig. 6C, D),** which was further increased if MCC950 was included (**Fig. S5A**). But the aggregation phenotype in response to nigericin was completely abrogated. These results indicate that ASC, potentially by cross-linking membrane-associated NLRP3, contributes to the TGN aggregation. Additionally, we further observed Golgi aggregation as the dominant feature in LPS- and nigericin-treated human primary macrophages that also contain all the NLRP3 inflammasome components (**Fig. S5B, C**).

Together, these results highlighted that TGN aggregation but not dispersion is a feature of NLRP3 membrane-bound dependent specks in fully inflammasome competent human immune cells. Considering this and our previous data we propose the following NLRP3 mechanisms of activation: 1) the NLRP3 double-ring cage resides at the TGN and upon K^+^-dependent or imiquimod stimulation a mild TGN dispersion is triggered. This allows NLRP3 to travel to endosomal-like structures which in turn culminates in TGN aggregation - only occurring if ASC is in the system - and further MTOC nucleation, resulting in the TGN/MTOC-dependent NLRP3 inflammasome activation; 2) smaller NLRP3 species such as monomers, cannot bind membranes and reside in the cytosol. These species are only able to respond to K^+^-dependent stimulation which triggers a strong TGN dispersion. Then, NLRP3 oligomerizes independently of membrane association, culminating in TGN/MTOC-independent NLRP3 activation. Interestingly, despite the presence of ASC in these cells, TGN aggregation does not occur. This suggests that both, membrane-associated NLRP3 and ASC are upstream of TGN aggregation (**Fig. 7**).

**Figure 7.**
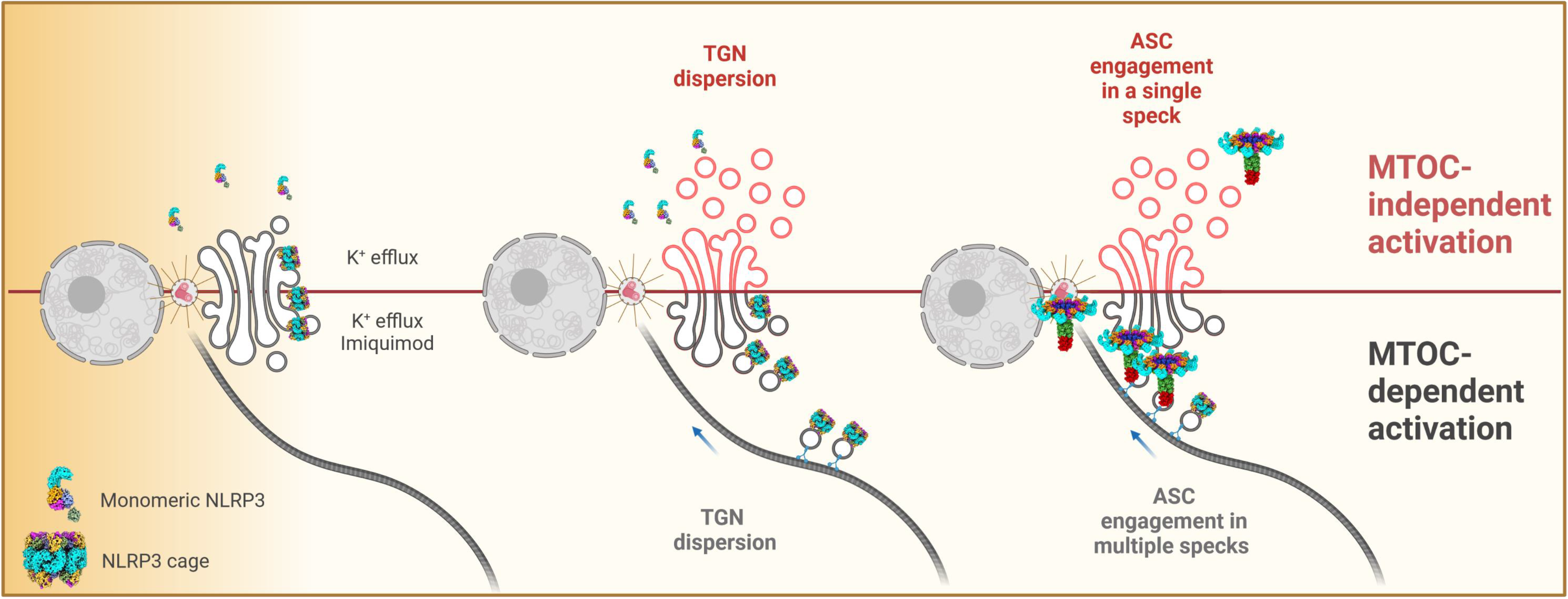
Proposed mechanism of TGN/MTOC-dependent and -independent NLRP3 inflammasome activation. The NLRP3 double-ring cage resides at the TGN and upon K^+^-dependent or imiquimod stimulation a mild TGN dispersion is triggered. This allows NLRP3 to travel to endosomal-like structures which in turn culminate in TGN aggregation - only occurring if ASC is in the system - and further MTOC nucleation, resulting in a TGN/MTOC-dependent NLRP3 inflammasome activation. However, smaller NLRP3 species, such as monomers unable to bind membranes, locate at the cytosol and are only able to respond to K^+^-dependent stimulation which triggers a strong TGN dispersion. Then, NLRP3 oligomerizes independently of membrane association and induces the TGN/MTOC-independent NLRP3 activation.

## Discussion

Over the years, NLRP3 activation has emerged to involve a staggering number of factors including protein conformation, modification, subcellular localization and trafficking. Our combined results establish new insights regarding the links between structural arrangement, localization and cell biology behind NLRP3 inflammasome activation. We here observed that the NLRP3 membrane-bound state is essential to locate NLRP3 in the right place within the cell to undergo activation; and this role seems to be orchestrated by the linker between the PYD and the NACHT domain encoded by the exon 3. Although this study has been performed by mainly using human THP-1 macrophage-like cells as a model, and further expansion upon the primary macrophage data we present will be helpful, we provide experimental evidence for the existence of parallel TGN/MTOC-dependent and -independent NLRP3 inflammasome activation pathways. In these, different states of the sensor protein (from monomers to cages) play a role in triggering an inflammatory response. Our study connects to ground-breaking structural work that has recently propelled our understanding of NLRP3 activation, while also opening up the discussion of whether different NLRP3 states from monomers to oligomers could all form active NLRP3 inflammasomes. However, mechanistic details, cell biological demonstration and the respective relevance for the ability of cells to respond to insults have lagged behind.

Here, we confirm that lipid-binding decameric human NLRP3 cages can follow the already described canonical murine TGN/MTOC-dependent NLRP3 activation pathway ^13, 24, 25^. Moreover, we demonstrate that smaller NLRP3 species incapable of organizing into decamers can engage a separate, TGN/MTOC-independent NLRP3 activation pathway. For WT NLRP3 both pathways can co-exist even within the same cell. Our work thus reconciles previously conflicting or ambiguous findings into a unified mechanism (**Fig. 7**). This proposed mechanism relates to initiation via both, TGN/MTOC/lipid binding and (presumably) cytosolic sites, either by NLRP3 oligomers or monomers, and leading to TGN aggregation or dispersion. These findings, as well as the role of NEK7 and the mechanism of CAPS *NLRP3* variants, which our two-pathway mechanism can accommodate and is extendable to, will be discussed in the following.

### Localization at steady-state

For instance, we observed that at basal conditions, NLRP3 located in heavy and light membranes as well as in the cytosol as previously reported ^20, 34^. What is noteworthy – and data from overexpression systems alluded to this – is that at endogenous expression levels in macrophage-like cells, other NLRP3 inflammasome components considered critical for full inflammasome execution – namely ASC, NEK7, pro-caspase-1 and pro-IL-1β – all reside in both membrane compartments and the cytosol before activation, albeit at slightly different proportions. These results suggest that, in principle, both locations are competent at executing full NLRP3 activation and support parallel TGN/MTOC-dependent and independent NLRP3 inflammasome pathways to coexist in human macrophage-like cells.

### NLRP3 structure: putative role of cages vs monomers in the TGN/MTOC-dependent and - independent NLRP3 activation pathways

In regard to NLRP3 structure, our work provides further proof of the notion that functional entities in NLRP3 are organized by exons: for example, the polybasic region (131-147) is split over two exons, exon 3 and exon 4. The deletion of the PYD-linker showed a human NLRP3 forming a hexamer ^24, 36^ but our deletion of only the linker sequence encoded by exon 3 led to smaller human NLRP3 species highly-enriched in monomers. In this sense, human NLRP3 linker could be primarily responsible for conferring the flexibility that allows two pre-requisites for appropriate stimulus-dependent activation: firstly, the PYD domains to be buried in the oligomer ^13, 14^; and secondly, to locate and orient the polybasic regions at the same interface for NLRP3 membrane-bound status. This notion is supported by the emergence of larger-than- monomer NLRP3 species solely by the presence of a six amino acids (GSGAGG) short flexible linker instead of the complete 40 amino acids encoded by the exon 3.

Earlier work proposed NLRP3 to exist from monomers to oligomers in a stable equilibrium ^13^, albeit it is unclear how, under cellular conditions, NLRP3 is sorted between cytosol and TGN. Our study clearly demonstrated the existence of two parallel NLRP3 activation pathways involving both monomers and cages: a TGN/MTOC-dependent pathway that could be modulated by the oligomer status of NLRP3, and a TGN/MTOC-independent one that appears governed by NLRP3 in the monomeric state. Consistently, the mostly monomeric Δexon3 NLRP3 construct, displays fully functional inflammasome properties upon K^+^-dependent stimulation independently of TGN/MTOC binding. In addition, endogenous skipping of exon 3 did not block K^+^-dependent NLRP3 inflammasome activation. A plausible explanation is that we suppressed the cage formation shifting the NLRP3 equilibrium to smaller species, consequently triggering the TGN/MTOC-independent pathway.

Importantly, the disruption of the microtubule system, a feature critical for the TGN/MTOC-dependent pathway in murine cells ^23^, in human cells failed to abolish the NLRP3 inflammasome activation associated to K^+^-dependent stimulation, further proving the existence of a TGN/MTOC-independent pathway. Interestingly, our results reconcile conflicting reports on NLRP3 membrane dependency due to the clear visualization of NLRP3 inflammasome specks that can sometimes be MTOC-associated or not. Additionally, we could also explain why an LRR-deleted NLRP3, which cannot form the double-ring cage and hence should exist in monomers, may form an active inflammasome when overexpressed ^19^, namely by the cytosolic fraction of this protein responding.

Strikingly, NLRP3 activation upon the K^+^-independent stimulus imiquimod showed a different scenario: Here, NLRP3 displayed a strong TGN localization and the addition of microtubule disrupting agents strongly dampened the inflammasome activation associated with imiquimod. Furthermore, the Δexon3 NLRP3 construct, unable to form cages, to bind TGN and to form MTOC-associated specks, did not respond to imiquimod stimulation. These results suggest that NLRP3 binding to membranes and residing with the MTOC system is critical for imiquimod response.

Regarding the physiological role of NLRP3 cages residing in the Golgi area, our live cell imaging experiments suggest that an increased concentration of NLRP3 located in this defined area triggers a more efficient and rapid inflammasome response which manifests itself in earlier ASC specks formation, IL-1β release, and pyroptotic cell death. This may be a mechanism to ensure rapid NLRP3 activation under suboptimal stimulation conditions.

### Human NLRP3 and PtdIns4P binding

Whilst NLRP3 sorting to membranes is unequivocal, our results demonstrate clear differences between mouse and human NLRP3 in terms of lipid affinity and membrane dependency. While human decameric NLRP3 only bound PA and PtdIns (3,4,5)-P_3_, mouse Nlrp3 appears more promiscuous and bound PA and all phosphorylated PtdIns ^13, 24^. This may explain the greater importance of membrane-associated NLRP3 activation in murine systems. PA presents a critical role in the Golgi structure and in the regulation of vesicle trafficking, and has been suggested as the source of diacylglycerol (DAG), a product of PA phosphatase and a second messenger for activation of protein kinase D, a key kinase involved in the NLRP3 activation pathway ^21, 37–40^. PtdIns (3,4,5)-P_3_ is exclusively located in the plasma membrane and is the target lipid of BTK, another kinase involved in NLRP3 activation ^26, 33, 41^, presumably only in the TGN/MTOC pathway consistent with the partial effect of BTK ablation on IL-1β release. We were not able to detect human NLRP3 directly binding PtdIns4P in the lipid strips, although there was detectable recruitment in PtdIns4P-rich structures – albeit not necessarily by direct interaction. PtdIns4P production is highly increased upon different NLRP3 inflammasome activators as a stress response mechanism employed by cells and PtdIns4P has been described for endosomal or lysosomal membranes ^22, 35^, whose damage was shown to contribute to NLRP3 inflammasome activation ^42^. We observed inactive NLRP3 recruited to PtdIns4P-enriched membranes in cells but upon nigericin stimulation this colocalization was reduced. Additionally, NLRP3 carrying the D305N mutation did not colocalize with PtdIns4P, albeit the recruitment was recovered in the presence of MCC950, an inhibitor enforcing cage structure ^14^. Thus, although human NLRP3 is visualized in PtdIns4P-rich membranes in the inactive state, presumably active or non-decameric NLRP3 does not preferentially reside with PtdIns4P. Furthermore, the Nlrp3 KKKK motif was directly linked to PtdIns4P binding and when mutated to AAAA, Nlrp3 activation was dampened in mice ^20^. However, we observed that the human NLRP3 KMKK motif mutated to AMAA responded to nigericin activation ^10^, suggesting a lower importance for this motif in human NLRP3. In addition, the auto-active phenotype of NLRP3 in the context of CAPS mutations D305N or L307P was still preserved when combined with the KMKK>AMAA polybasic mutation ^43^. Thus, PtdIns4P production may contribute to trigger NLRP3 activation but seems not essential for the switch to the actual human NLRP3 activation platform.

### TGN dispersion *vs* aggregation

Although initially opposing dispersion *vs* aggregation phenotypes observed in our experiments may seem confusing, our data suggest that both processes appear to coexist and may facilitate NLRP3 inflammasome activation. Upon nigericin stimulation, where both TGN/MTOC-dependent and -independent pathways occur, the two phenomena dispersion and aggregation were visible in WT NLRP3 reconstituted THP-1 cells as well as in human primary macrophages. The fate of the TGN probably correlates with which type of pathway dominated in each cell. However, the Δexon3 NLRP3 construct, unable to bind membranes, only led to dispersion, via the TGN/MTOC-independent NLRP3 activation pathway. Conversely, imiquimod stimulation which only triggers the NLRP3 TGN/MTOC-dependent activation pathway, showed only TGN aggregation. Interestingly, the aggregation phenomenon occurred only in the presence of ASC. TGN aggregation feature may have been overlooked so far due to most studies on TGN dispersion have been performed in non-macrophage cells that lack naturally the inflammasome components or specifically in an *ASC* KO background ^20, 22, 23, 35^. We associate aggregation more with the TGN/MTOC-dependent NLRP3 activation pathway because aggregation is a direct consequence of NLRP3 located in the TGN/MTOC membranes, while dispersion seems linked to the TGN/MTOC-independent one. Interestingly, two other groups indirectly support our proposed two-pathway model contributing to maximal NLRP3 activation: Both Bali Lee *et al*., and Schmacke *et al*., were able to potentiate imiquimod (i.e. K^+^-independent but TGN/MTOC -dependent) activity by adding either a TGN dispersing agent (monensin) ^35^ or completely K^+^-free buffer (forcing K^+^ efflux) ^34^, thereby enrolling the additional TGN/MTOC-independent TGN aggregating pathway. These results are in agreement with our observations and support the parallel activation of the NLRP3 TGN/MTOC-dependent and -independent pathways.

### The potential role of NEK7

The two-pathway model also helps accommodate the controversial role of NEK7 in NLRP3 activation. Cryo-electron microscopy studies have shown both the structure of the inactive human monomeric NLRP3 in complex with NEK7 ^17^ as well as the structure of the active human NLRP3 disc with 1:1 NLRP3-NEK7 complexes ^25^. However, NEK7 was unable to bind murine NLRP3 in the inactive cage conformation as the interaction sites are buried in the NLRP3 structure ^13, 14^. In line with these structural studies, NEK7 should also be able to bind monomeric Δexon3 NLRP3 species because the amino acids responsible for NEK7 interaction should be available. Consistent with our results, in the same way that Δexon3 NLRP3 fails to respond to imiquimod, NEK7 KO cells fail too as shown recently by Schmacke *et al* ^34^. By analogy, we anticipate only the human cage NLRP3 to be dependent on the NEK7 action. In this sense, we favour the proposed role of NEK7 in disrupting the NLRP3 cage into two halves to allow an active disc formation during the activation state ^25^. However, in the absence of NEK7, monomeric WT NLRP3 could still respond to nigericin stimulation via the NLRP3 TGN/MTOC-independent pathway similarly to Δexon3 NLRP3.

### Autoinflammatory CAPS: a potential TGN/MTOC-independent activation phenotype

The clear involvement of the NLRP3 inflammasome in promoting inflammation in CAPS patients is at odds with the uncertainties about how CAPS mutated NLRP3 actually works molecularly. Normally, cells in which NLRP3 CAPS mutants have been studied ectopically present a pattern of NLRP3 foci distributed around the cytosol which point to an TGN/MTOC-independent activation pathway ^9, 20, 21, 43^. This is consistent with the proposed observation that the constitutively active NLRP3 disease mutants could bypass TGN recruitment also in the presence of the mutant L351P carrying the extra KKKK>AAAA mutation because they were all able to aggregate without exogenous stimulation ^20^. Moreover, an LRR-deleted NLRP3 unable to organise in double-ring cages and further carrying the R260W or T348M CAPS mutations still supported constitutive activity ^19^. Additionally, the NLRP3 L353P mutant efficiently responded to cold stimulation in the absence of NEK7, further consistent with the TGN/MTOC-independent pathway as discussed above ^43^.

From a therapeutic angle, colchicine treatments have been applied to relieve the inflammation symptoms associated with NLRP3 hyperactivation such as gout flares ^44^. For CAPS treatment, colchicine has not been extensively used, but the few studies performed on these patients showed only a partial or no response ^45^, indicating poor efficacy. This clinical observation squares well with the dominance of an MTOC-independent activation pathway also in CAPS. Elucidating to which extent TGN/MTOC-independent pathway could govern the mechanism of NLRP3 activation in CAPS-affected patients and other non-genetic NLRP3-related pathologies might lead to new alternatives and more efficient methods to block inflammation in each setting.

## Acknowledgements

We thank Gloria López-Castejón and Fátima Martin-Sánchez (Institute of Immunology and Inflammation, University of Manchester) for providing THP-1 cells used in this paper, our students Kim Hebel and Garvit Sharma for their help with ELISAs, Kay Oliver Schink (University of Oslo) for pENTR20-mNG plasmid and for sharing very useful Fiji scripts for the analysis of our microscopy data, Michael Schindler (Molecular Virology, University of Tübingen) for the TGN46 plasmid, Vinicius Nunez Cordero Leal for PBMC isolation, Markus W. Löffker for biobank support and acquiring blood samples, Sarah-Maria Trenz and Samuel Wagner (Cellular and Molecular Microbiology, University Tübingen) for their support with cell fractionation, and Kia Wee Tan (Uppsala University) for his useful advice and comments on cellular trafficking. The study was supported by the grants from the internal support program of the Medical Faculty, University of Tübingen, Fortüne-Antrag Nr. 2615-0-0 and Nr. 3023-0-0 (to M.M.-T and A.T.-A), and the DFG (German Research Foundation) Clusters of excellence “iFIT-Image Guided and Functionally Instructed Tumor Therapies” (EXC-2180, to A.N.R.W and A.T.-A) and “CMFI-Controlling Microbes to Fight Infections” (EXC-2124, to J.G and A.T-A). M.G. is funded under Germany’s Excellence Strategy–EXC2151–390873048 and by grant of the DFG (GE 976/16-1).

## Author contributions

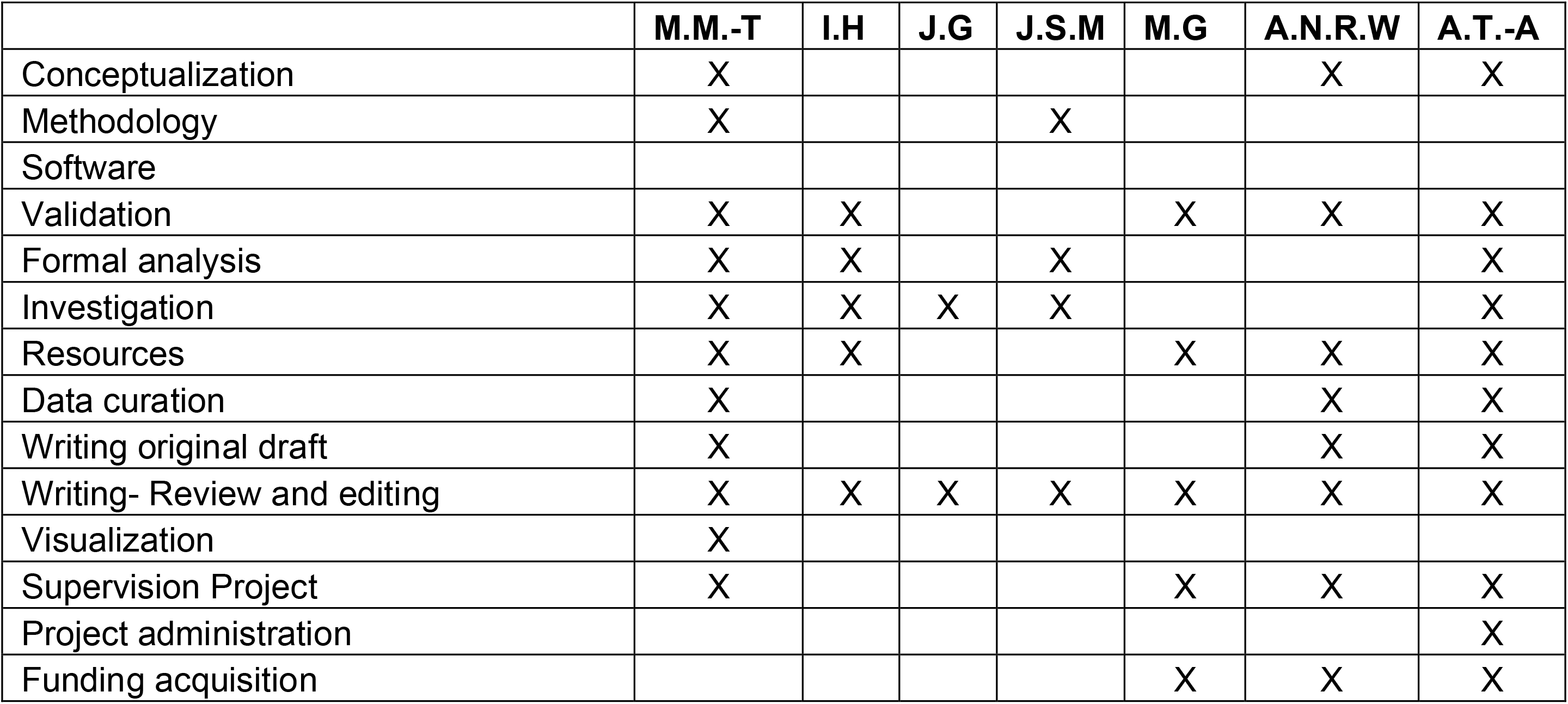

## Methods

### Chemicals and reagents

Phorbol 12-myristate 13-acetate (PMA), LPS ultrapure from *Escherichia coli* K12 (LPS-EK, tlr-peklps), nigericin sodium salt (tlr-nig), imiquimod (tlrl-imq) and monosodium urate crystals (MSU) were obtained from InvivoGen. MCC950 was purchased from MedChemExpress, LukAB was a kind gift from C. Wolf ^33^. Colchicine, Nocodazole and Ciliobrevin D were acquired from Sigma Aldrich, Ricolinostat from SelleckChem.

### Plasmids and cloning procedures

The different ENTRY plasmids of human NLRP3 were generated by PCR from an NLRP3 template (Uniprot #Q96P20) and cloned into pENTR20-mNG (kind gift from Kay Oliver Schink, University of Oslo). Sequencing was performed to confirm correct amplification and the absence of unwanted mutations. pTGN46-mScarlet2 was a kind gift from Michael Schindler (Molecular Virology, University of Tübingen) in which mScarlet2 was replaced with mCherry. GFP-2xSidM (Addgene plasmid #51472) in which GFP was exchanged with mCherry. All other plasmids were generated using standard molecular cloning techniques. Detailed cloning procedures are available from the authors.

### Cell culture, treatments, and transfection

HeLa cells were cultured in complete DMEM (4.5 g l^-1^ glucose, containing 10% heat-inactivated fetal bovine serum, 2 mM L-Glutamine, 100 U/mL penicillin, 100 µg/mL streptomycin) and sub-cultured every 2-3 days. For some experiments, HeLa cells were transfected with 2xSidM or TGN46 plasmids using Lipofectamine 2000 according to the manufactures’ instructions. After 24 h of transfection the experiment was performed.

THP-1 WT cells (kind gift from Thomas Zillinger, Bonn, Germany) were maintained in complete RPMI 1640 (containing 10% FBS, 25 mM HEPES, 2 mM L-Glutamine, 100 U/mL penicillin/100 µg/mL streptomycin) and sub-cultured every 3 days. THP-1 Null 2 (InvivoGen), THP-1 Null 2 *NLRP3* KO (InvivoGen) and THP-1 Null 1 ASC KD (InvivoGen) were maintained in complete RPMI 1640 (containing 10% heat-inactivated fetal bovine serum, 25 mM HEPES, 2 mM L-Glutamine, 100 U/mL penicillin,100 µg/mL streptomycin, 100 µg/mL Normocin, and 100 µg/mL of Zeocin) and sub-cultured every 3 days. All cell lines were cultured at 37 °C in 5% CO_2_ and regularly checked by PCR to avoid *Mycoplasma* contamination.

For assessing NLRP3 activation, HeLa cells were stimulated with 10 µM nigericin for 1 h in the presence or absence of 10 µM MCC950 in OptiMEM. For inflammasome stimulation in THP-1 cells, they were first differentiated with 100 ng/mL of PMA overnight, rested for 2 days, then primed with 10 ng/mL of LPS-EK for 4 h and stimulated or not with 10 µM nigericin, 100 µM imiquimod, 250 µg/mL MSU or 5 µM LukAB at the indicated times. For inflammasome inhibition 10 µM MCC950 was added into the cultures 30 min before and during inflammasome activation. Where indicated, 40 mM KCl was added to the culture media in order to prevent potassium efflux during inflammasome activation.

For pharmacological inhibition, LPS-primed THP-1 cells were pretreated for 1 h with the HDAC6 inhibitor Ricolinostat (10 µM). Dynein-dependent microtubule transport was inhibited using Ciliobrevin D (10 µM). For microtubule polymerization disruption, nocodazole (2.5-10 µM) or colchicine (10 µM) were used. After 1 h, cells were treated for inflammasome activation.

### Generation of stable cell lines

Third-generation lentivirus was generated using procedures and plasmids as previously described ^46^. Briefly, NLRP3 WT, D305N, Δexon3 and AMAA constructs all C-terminally mNG-tagged and previously generated as Gateway ENTRY plasmids were transferred by Gateway LR recombination into lentiviral destination vectors ^47^ (Gateway-enabled vectors derived from pCDH-EF1a-MCS-IRES-BLAST, (SystemBiosciences). VSV-G pseudotyped lentiviral particles were packaged using a third-generation packaging system (Addgene plasmids 12251, 12253, 12259).

For the generation of lentiviral particles, 0.5 µg pCMV-VSVG, 1 µg pMDLg/pRRE and 1 µg pRSV-REV plus 2 µg of the gene of interest expressing plasmid were transfected into HEK293T cells using Lipofectamine 2000 on a final volume of 1 mL and following the manufacturer’s instructions. The day before transfection, 1×10^6^ of freshly thawed HEK293T were seeded in a 6-well plate in complete DMEM (4.5 g l^-1^ glucose, 10% FBS, 1% L-Glutamine, 1% P/S). After 72 h, the supernatants were collected and filtered using 0.45 µm Millex-HV Syringe Filters and directly used for transduction or kept at −80 °C. HeLa cells were infected with a 25% (v/v) of lentivirus-containing media and incubated for 3-5 days. To improve the efficiency of transduction in THP-1 NLRP3 KO, both cells and 50% of lentivirus-contained supernatant were centrifuged at 1000 × *g* at 30°C for 1 h. Then, the THP-1 cells were resuspended up and down gently and seeded and incubated for 3-5 days. To obtain stable expressing populations, HeLa and THP-1 cells were subjected to antibiotic selection using blasticidin at 5 µg/mL.

### hMDMs isolation, macrophage differentiation and inflammasome activation

All healthy donors included in this study provided their written informed consent before participation. Approval for use of biomaterials was obtained by the local ethics committee of the Medical Faculty Tübingen in accordance with the principles laid down in the Declaration of Helsinki as well as applicable laws and regulations.

Peripheral blood mononuclear cells (PBMCs) were isolated from whole blood by density centrifugation with Ficoll (1.077 g/mL, Sigma, 10771). Briefly, EDTA-anticoagulated blood samples were diluted 1:1 with PBS (Thermo Fisher, 14190-169) and subsequently added onto Ficoll and centrifuge at 500 × *g* for 25 min at room temperature without brake. Afterwards, the PBMC layer was carefully collected and transferred into another tube and washed twice with PBS. In order to isolate the monocytes from the PBMCs portion, 30 × 10^6^ cells were seeded in 10 cm culture-treated dishes (Greiner bio-one) in 10 mL of Monocyte Attachment Medium (Promocell, Heidelberg, Germany) to promote monocyte adhesion. After 1.5 h of incubation at 37 °C and 5% CO_2_, the supernatant was aspirated from the dishes and the dished washed 3 times with PBS to eliminate non-adherent cells. Then, complete RPMI medium (10% FBS, 1% L-glutamine, 1%P/S) was added and monocytes were differentiated into macrophages with 50 ng/mL of hGM-CSF (Sargramostin, Leukine/Sanofi). The cells were incubated for 7 days refreshing the culture at day 4 with 5 mL of complete RPMI medium and 50 ng/mL of hGM-CSF. Afterwards, the macrophages were washed twice with PBS and subsequently detached adding 10 mL of Macrophages Detachment Medium (Promocell, Heidelberg, Germany). The dishes were incubated for 40 min at 4 °C and 20 min at room temperature, and then, 10 mL of complete RPMI medium were added to finally detach the cells carefully pipetting up and down. The collected cells were washed twice complete RPMI medium, and the cell concentration determined.

The macrophages obtained were seeded in coverslips at 0.3 × 10^6^ cells/well in 500 µL of complete RPMI medium and incubated for 24 h at 37 °C and 5% CO2 to facilitate their adhesion. For NLRP3 inflammasome activation, hMDMs were primed with 100 ng/mL of LPS-EK for 4 h and stimulated or not with 10 µM nigericin for 1 h.

### Cloning, expression, and purification of human NLRP3

Full length, human NLRP3 (3-1036, UniProt accession code Q96P20) wildtype or mutants with the linker region (exon 3 sequence) deleted (Δ95-134) or deleted and reconstituted with a small, flexible linker (Δ95-134 + linker (GSGAGG)), codon-optimized for *Spodoptera frugiperda*, were cloned in an in-house modified pACE-Bac1 vector containing a N-terminal MBP-tag, followed by a Tobacco etch virus (TEV) protease cleavage site.

For recombinant protein expression of the different NLRP3 proteins, 500 mL of *Sf9* insect cells were infected with 3 % v/v viral stock of the second virus passage. Expression cultures were incubated for 48 h at 27 °C and 80 rpm, and were subsequently harvested by centrifugation at 2000 rpm for 20 min. Cell pellets were washed with PBS and either subsequently used for protein purification or flash-frozen in liquid nitrogen for storage at -80°C. For protein purification, cell pellets were solubilized in lysis buffer (50 mM HEPES pH 7.5, 150 mM NaCl, 0.5 mM TCEP, 10 mM MgCl_2,_ 1 mM ADP), supplemented with 1 mM phenylmethylsulfonyl-fluoride (PMSF), followed by sonication (10 sec on, 5 sec off for 4 min at 40% intensity) on ice. Cell lysates were centrifuged at 25000 rpm and 4 °C for 1 h and the supernatants were filtered with a 45 µm syringe filter, prior to application onto a lysis-buffer equilibrated 5 mL MBP-trap column (GE Healthcare) attached to an ÄKTA-Start FPLC system. The column was subsequently washed with 10 CVs of lysis buffer, followed by elution of the proteins with 5 CVs of lysis buffer supplemented with 15 mM maltose. Proteins were further purified by size exclusion chromatography (SEC) on a Superose 6 increase 10/300 GL column (GE Healthcare) equilibrated with lysis buffer. Elution fractions were analyzed by SDS-PAGE and negative stain electron microscopy (EM). Proteins used for the lipid strip binding assays were purified as described before, except for the supply of 10 µM MCC950 in the expression cultures and purification buffers.

### Negative stain EM

For negative stain EM, 4 µL of the indicated SEC elution fractions were applied onto glow-discharged carbon-coated copper grids (200 nm mesh, PLANO). Samples were incubated on the grid for 1 min, before excess sample was blotted away with a filter paper. Grids were subsequently washed by dipping the sample side into three individual 20 µ L drops of lysis buffer, alternated by a blotting step in between. Samples were negative stained by dipping the sample side into a 20 µL drop of 2 % uranyl acetate and incubation for 30 s, before excess stain was blotted away with a filter paper. Samples were air-dried and subsequently imaged with a Jeol JEM-2200FS transmission electron microscope operated at 200 kV.

### SEC-coupled Multi-Angle Light Scattering (SEC-MALS)

For SEC-MALS of the MBP-NLRP3 (Δ95-134) protein, elution fractions (C7-D4) corresponding to the second peak of the SEC purification, were pooled and concentrated to 1.3 mg/ml). The concentrated protein sample was centrifuged at 10,000 × *g* and 4°C for 10 min before the supernatant was loaded onto a lysis buffer equilibrated Superose 6 increase 10/300 GL column (GE Healthcare), connected to a 1260 Bioinert Infinity LC system and equipped with a multi-angle light scattering detector (miniDawn 3141MD3, Wyatt) and a refractive index detector (Optilab rEX 650, Wyatt). Data were collected every 0.5 s applying a flow-rate of 0.5 ml min^-1^. Data analysis was performed using the Astra 8 software (Wyatt).

### Immunofluorescence analysis

Cells were imaged using a Plan-Apochromat 63×/1.4 oil objective on a Zeiss LSM800 confocal system equipped with an Airyscan module and controlled by the Zen blue software. In brief, cells plated in coverslips (12 mm) were fixed in 4 % PFA in PBS at 37 °C for 15 min, then washed in PBS 3 times with 5 min interval between washes. Afterwards, cells were permeabilized for 5 min in 0.05 % saponin in PBS for 5 min. Cells were then blocked in blocking buffer (2 % bovine serum albumin (BSA), 0.05 % saponin in PBS) for 1 h. Staining with primary antibodies (2 h to overnight) and secondary antibodies (1 h) was performed in blocking buffer. To remove unbound antibodies, in between washing steps were performed. Staining of the nuclei was performed with Hoechst 33342. The primary and secondary antibodies used for immunofluorescence in this study are listed in in Supplementary Table 1.

### Live Cell imaging

Live cell imaging experiments were performed in an LSM 800 (Zeiss) microscope equipped with a 63x 1.42 NA Objective (Zeiss). Environmental control was provided by the incubation chamber (controlled temperature, humidity, CO_2_), a heated stage (Heating insert P Lab-TekTM S1, Zeiss) and an objective heater (Zeiss). Briefly, PMA-differentiated THP-1 Null2 *NLRP3* KO cells reconstituted with WT- or Δexon3-mNG NLRP3 were seeded in 4-chamber glass-bottom dishes (Ibidi). After priming with LPS-EK for 4 h, cells were stimulated while imaging with nigericin (10 µM) or imiquimod (100 µM) in phenol red-deficient RPMI 1640 imaging media (Sigma). In order to stain cell death, SYTOX orange (Invitrogen, dilution 1/50000) was added to the media 10 min prior imaging. To stain the microtubule network, SiR-Tubulin incubation was performed for 4 h (Cytoskeleton, with 2 μM SiR-Tubulin and 10 μM verapamil). Videos were taken for 2 h at intervals of 48-120 s. Analysis of live cell imaging and generation of the different galleries were performed using Fiji. In brief, individual cells were manually tracked (10-20 cells approx. per condition and per experiment) and each frame was added to the ROI manager. Galleries of tracked cells were generated with the automated macro “MeasureTracks_Gallery” (https://github.com/koschink/Phafin2).

### Immunoblot

After treatments, cell culture supernatants and cell lysates were collected for immunoblot analysis. Cells were lysed for 30 min on ice cold 1× RIPA buffer (50 mM Trizma base pH 7.4, 150 mM NaCl, 1 mM EDTA, 1 % Triton X-100, 0.1 % SDS, 0.5 % sodium deoxycholate, 10 % glycerol) supplemented with one tablet of protease inhibitor cocktail (Roche) for every 10 mL of RIPA. Cell lysates were centrifuged at 16000 × *g* for 15 min at 4 °C. For the supernatants, the proteins were precipitated and concentrated using methanol–chloroform precipitation, and finally resuspended in RIPA buffer. The immunoblots were prepared using Tris-glycine SDS– PAGE and transferred to nitrocellulose membranes (Bio-Rad) by electroblotting. Membranes were blocked with Tris-buffered saline with 0.1 % Tween 20 (TBS-T) + 5 % BSA (blocking buffer) and probed overnight at 4 °C with primary antibody. After three washes with TBS-T, membranes were incubated with HRP-conjugated secondary antibody in blocking buffer for 1 h at room temperature. Membranes were again washed with TBS-T three times. Proteins were visualized using ECL substrate with chemiluminescent detection (LI-COR Odyssey). The primary and secondary antibodies used for immunoblotting in this study are listed in in Supplementary Table 1.

### Subcellular fractionation

THP-1 cells were homogenized using a 10 mL syringe and 27 G × 19 mm needles in homogenization buffer (0.25 M sucrose, 10 mM Tris HCl pH 7.5, 10 mM KCl, 1.5 mM MgCl _2_, protease inhibitor (Roche). Homogenized cells were centrifuged at 1000 × *g* for 5 min to remove the nucleus. The supernatant was centrifuged at 5000 × *g* for 10 min to obtain the heavy membrane fraction (pellet, P5) and the supernatant (S5) was centrifuged at 100000 × *g* for 20 min to separate the light membrane fraction (pellet, P100) from the cytosol + vesicle fraction (supernatant, S100). The samples were further analysed by immunoblot with equal amounts of protein loaded per lane.

### In vitro lipid strip assay

Lipid binding assay with purified human NLRP3 variants was performed using PIP strips (Echelon Biosciences) according to manufacturer instructions. In short, the PIP strip membranes were blocked using 3 % BSA in TBS-T for 1 h followed by incubation with 5 mL of WT or Δexon3 NLRP3 proteins (7.5 μg in total) diluted in 3 % BSA in TBS-T for 2 h. Next, the membranes were washed in TBS-T for 15 min and incubated with the primary antibody Cryo2 1:1000 in TBS-T 3 % BSA for 1 h. Then, the strips were washed in TBS-T for 15 min and incubated with the secondary antibody anti-mouse IgG-HRP (Promega, Cat. no: W4028) at 1:10000 dilution in TBS-T 3 % BSA for 1h. Then, the strips were washed in TBS-T for 30 min and the bound proteins were visualized using ECL substrate with chemiluminescent detection (LI-COR Odyssey). All steps were performed at room temperature. All samples were analysed at the same time under the same conditions.

### Cytokine measurement

Human IL-1β (Biolegend ELISA MAX Deluxe kit, 437015) and human TNF-α (Invitrogen, 88-7346-22) were measured by triplicate ELISA according to manufacturer’s instructions.

### Lactate dehydrogenase assay

To measure cell death, the LDH present in cell-free supernatants was detected in triplicates using the Cytotoxicity Detection kit (Roche) according to manufacturer instructions, the reaction was read in a Synergy Mx (BioTek) plate reader at 492 nm and corrected at 620 nm.

### SYTOX Orange assay

In order to assess cell death, the signal of SYTOX™ Orange Nucleic Acid Stain (Invitrogen, S11368) was measured in the plate reader in real time in living cells. In brief, THP-1 cells were prepared for inflammasome activation in 96-well black plates (Thermo Fisher, 165305) in the presence of SYTOX™ orange (dilution 1/50.0000). Kinetics was measured by detecting SYTOX signal every minute with a filter 530/25,590/35 using the Synergy 2 plate reader (BioTek) at 37 °C in 5 % CO_2._

### Caspase-1 activity

Staining of active caspase-1 was performed with carboxyfluorescein (FAM) labeled inhibitor of caspase-1 (FAM-FLICA 660™) kit (ImmunoChemistry Technologies) according to manufacturer’s protocol. After inflammasome activation, media was removed and cells were incubated for 1 h in complete RPMI with 1x FAM-FLICA. Cells were then washed in complete RPMI for another hour and then fixed in 4 % PFA as described in the immunofluorescence section.

### Design of antisense oligonucleotides for exon 3 skipping

Antisense oligonucleotides targeting the exon 3 were designed following protocols previously described ^48^. Briefly, potential targets sites for efficient exon skipping were identified using the server Human Splicing Finder (Genomnis SAS Company, France) ^49^. Two potential regions were selected based on the prediction of possible exonic splice enhancer sites (ESEs). The predicted mRNA structure by mFoldWeb Server ^50^ showed that one of those regions was partially open, a necessary requisite for a potential target region. The final designed oligonucleotide was named AON exon3 (*5’* CACTCCTCTTCAATGCTGTCTTCCT *3’*).

### RNA isolation, NLRP3 qPCR

RNA was isolated using the RNeasy kit from Qiagen, according to the manufacturer’s instructions and quantified on Nanodrop 1000 (Thermo Fisher Scientific). Reverse transcription of RNA (800-1000 ng) to generate cDNA was performed using High*-*Capacity cDNA Reverse Transcription Kit (Thermo Fisher Scientific) following manufacturer’s instructions. In order to assess the relative expression of the exon 3 (skipped mRNA) and exon 7 (total mRNA), cDNA abundance was measured by qPCR in a QuantStudio6 cycler using TaqMan^TM^ system (Thermo Fisher Scientific) following manufacturer’s instructions. The TaqMan™ Gene Expression Assays used were the following: for TBP (housekeeper) Hs00427620_m1, for NLRP3 exon 2-3, ID Hs00918082_m1; for NLRP3 exon 6-10, ID Hs00918080_g1. Analysis was performed using QuantStudio Real-Time-PCR software version 1.3 (Thermo Fisher Scientific).

### Antisense oligonucleotides and exon skipping protocol

Control antisense oligonucleotide (Control AON), AON targeting exon 5 ^29^, AON targeting exon 3 and Endoporter were obtained from gene-tools.com. THP-1 WT cells were treated overnight with PMA (100 ng/mL) for differentiation. Media was changed the day after for complete RPMI. After 48 h, transfection was performed with premixed morpholinos and Endoporter to a final concentration of 10 µM and 6 µM, respectively. Of note, morpholinos were heated up to 65 °C in order to recover full activity before use. 40 h after transfection, cells were treated for inflammasome activation.

### Image analysis

Image processing and quantification was performed in Fiji software. For colocalization analysis, Pearson’s correlation with Costes automatic threshold was applied by using JacoP Plugin. Quantification of NLRP3 specks associated with TGN38 was performed by segmentation of NLRP3 specks with the automatic threshold Renyi’s entropy and regions of interest (ROIs) were generated with the plugin Analyze particles. Defined ROIs were superimposed onto the channel of interest (TGN38) and mean intensity was measured. For determining membrane aggregation and dispersion, segmentation of TGN38 or RCAS1 signal was performed by using the threshold Triangle. ROIs were defined with the plugin Analyze particles and the mean intensity or the area was measured. Quantification of percentage of ASC specks was performed by segmentation of ASC and nuclei signal by thresholding. Number of ASC specks and nuclei were determined by using the plugin Analyze particles and the ratio between them was calculated. The percentage of TGN and MTOC associated specks was performed manually in 3×3 TILES (63x). For live cell imaging experiments, timing of speck formation in MTOC-dependent or -independent specks was determined manually. The distance travelled by WT or Δexon3 specks was determined using Mtrack2 plugin from Fiji. For each experiment, images were randomly acquired with the same settings (laser power, detector gain) and below pixel saturation.

### Statistics

Experimental data were analysed using GraphPad Prism 8. Normal distribution in each group was analysed using the Shapiro-Wilk test first for the subsequent choice of the parametric (ANOVA when comparing multiple groups or Student’s t-test when comparing two groups for normally distributed data) or non-parametric (Mann-Whitney U) test as indicated in the figure legends. Data are always presented as mean ± SD, *p*-values (α=0.05) were calculated as indicated in the figure legends using GraphPad Prism and values <0.05 were considered statistically significant. For visual annotation of statistical significance in graphs, the following nomenclature was used: *p < 0.05, **p < 0.01, ***p < 0.001, **** < 0.0001, ns, not statistically significant.

## Supplemental information

**Figure S1.**
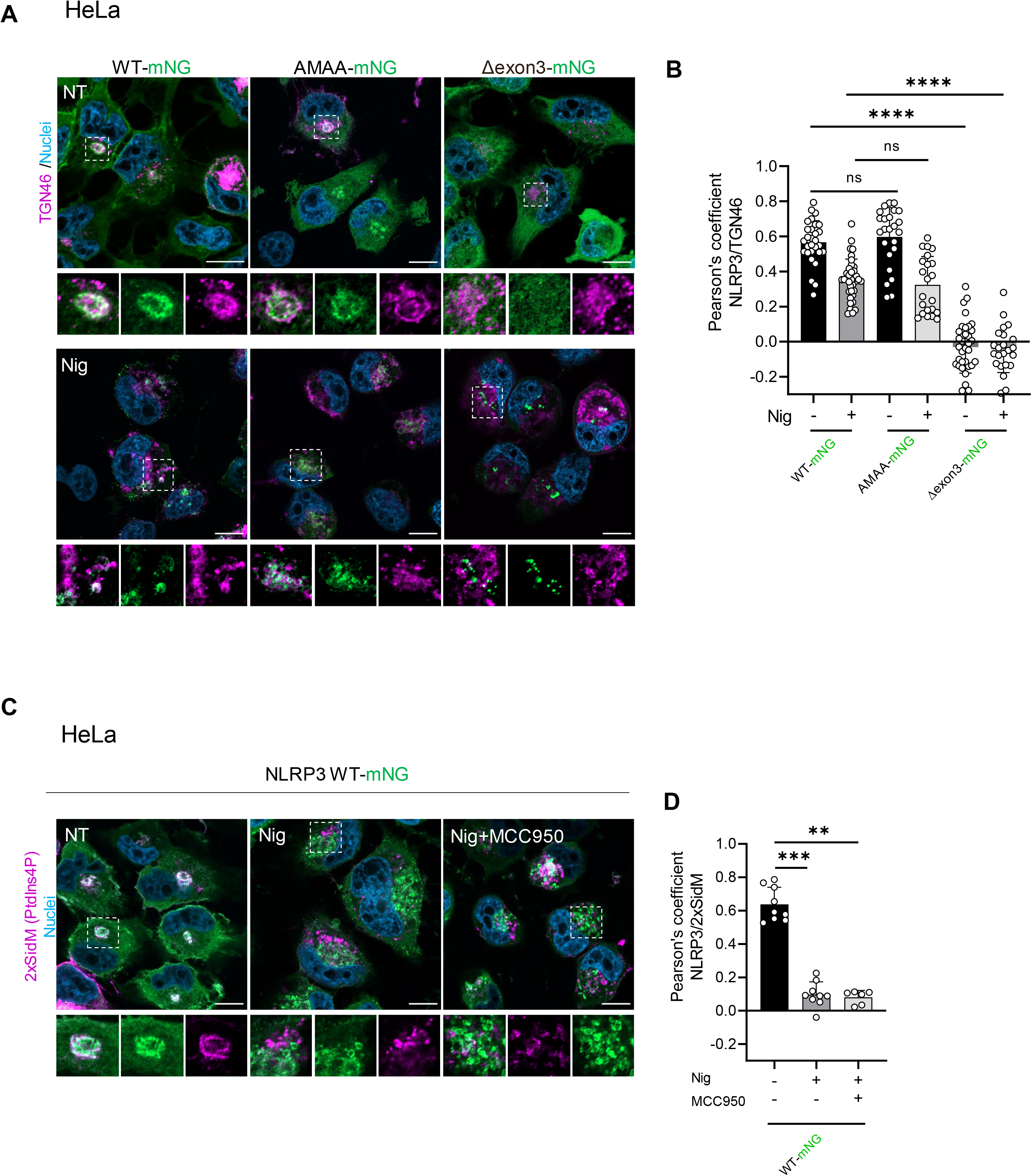
Human Δexon3 NLRP3 expressed in HeLa cells is not located in the TGN membranes in basal nor upon nigericin activation. **A)** Representative confocal microscopy images of HeLa cells stably expressing human NLRP3 WT, NLRP3 with the KMKK motif mutated to AMAA or NLRP3 Δexon3 all C-t tagged with mNG (green), and co-transfected with the TGN marker, TGN46-mCherry (magenta) and then treated or not with nigericin at 10 μM for 1 h. Nuclei were stained with Hoechst 33342 (blue). Scale bar 10 µm, N=2 independent experiments. **B)** Co-localization analysis of WT, AMAA or Δexon3 NLRP3 and TGN treated as in **A**. Values are represented by Pearsońs correlation coefficient with Costes automatic thresholding. Each bar represents the mean ± SD with each data point representing one cell from 2 independent experiments (around 20 cells per independent experiment). ****p≤0.0001; ns non-significant according to Kruskal-Wallis test with Dunńs multiple comparisons test. **C)** Representative confocal microscopy images of HeLa cells stably expressing human WT NLRP3-mNG (green) and transfected with the PtdIns4P sensor, 2xSidM-mCherry (magenta) treated or not with nigericin. Nuclei stained with Hoechst 33342 (blue). Scale bar 10 µm. N=3 independent experiments. **D)** Co-localization levels of WT NLRP3 and PtdIns4P of **C**. Each bar represents the mean ± SD with each data point representing one cell from 3 independent experiments (3 cells per experiment). ***p≤0.001; **p≤0.01 according to Kruskal-Wallis test with Dunńs multiple comparisons test.

**Figure S2.**
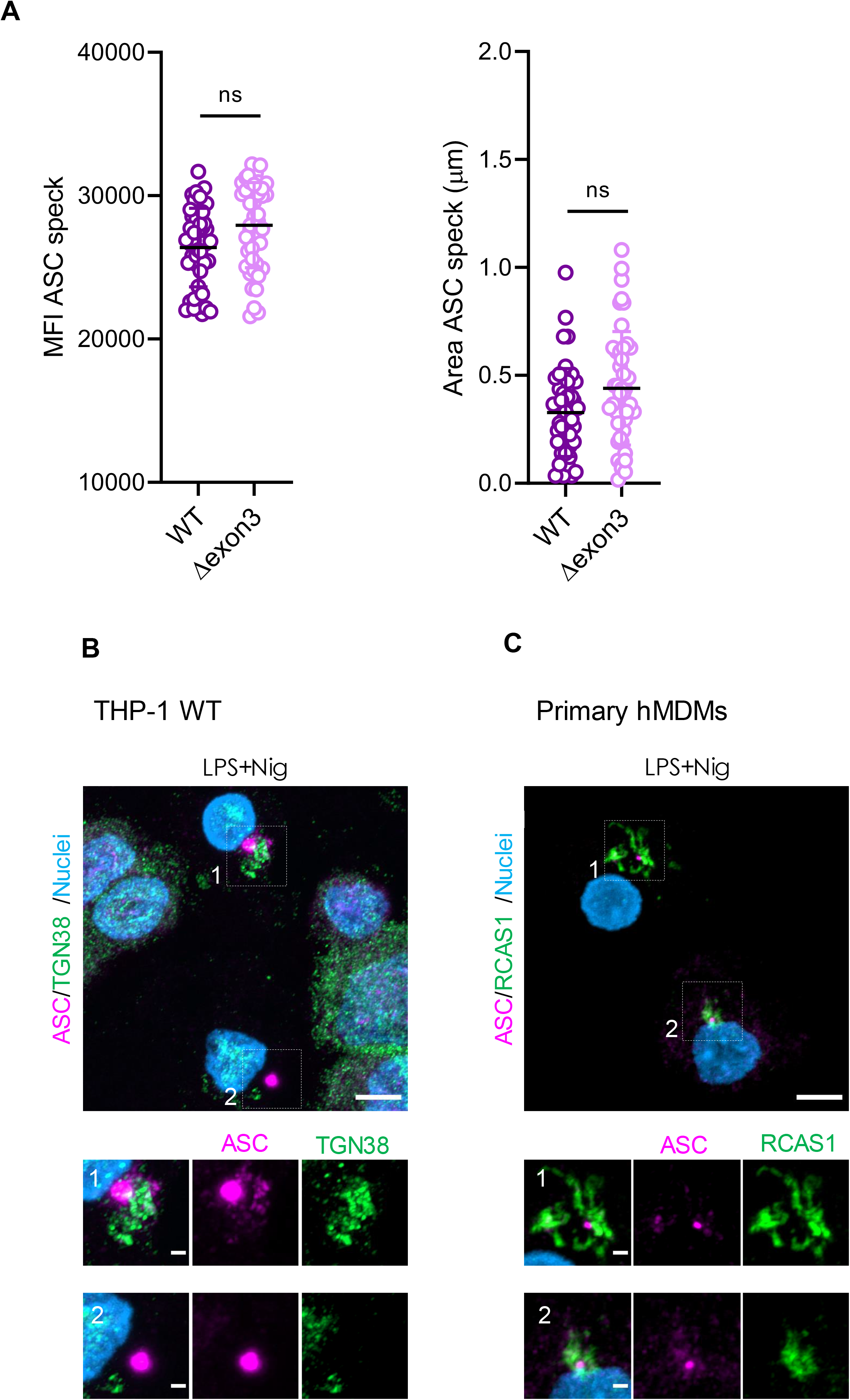
NLRP3-ASC platforms form TGN- and MTOC-independent specks in response to nigericin. **A)** Fluorescence intensity (left) and area (right) of the TGN-independent NLRP3-ASC platforms of the indicated LPS-primed and nigericin-stimulated THP-1 cell lines. Data are represented as mean ± SD and each data point represents one speck from 3 independent experiments. ns, non-significant according to according to Mann*–*Whitney U test. **B)** Representative immunofluorescence images of LPS-primed and nigericin-stimulated THP-1 cells with endogenous ASC stained using a specific ASC antibody (magenta) and TGN with TGN38 antibody (green). Nuclei were stained using Hoechst 33342 (blue). Scale bar 5 µm, zoom in 1 µm. N=1 independent experiment. **C)** Representative immunofluorescence images of LPS-primed and nigericin stimulated human primary macrophages with endogenous ASC stained using a specific ASC antibody (magenta) and Golgi membranes with RCAS antibody (green). Nuclei were stained using Hoechst 33342 (blue). Scale bar 5 µm, zoom in 1 µm. N=3 independent experiments.

**Figure S3.**
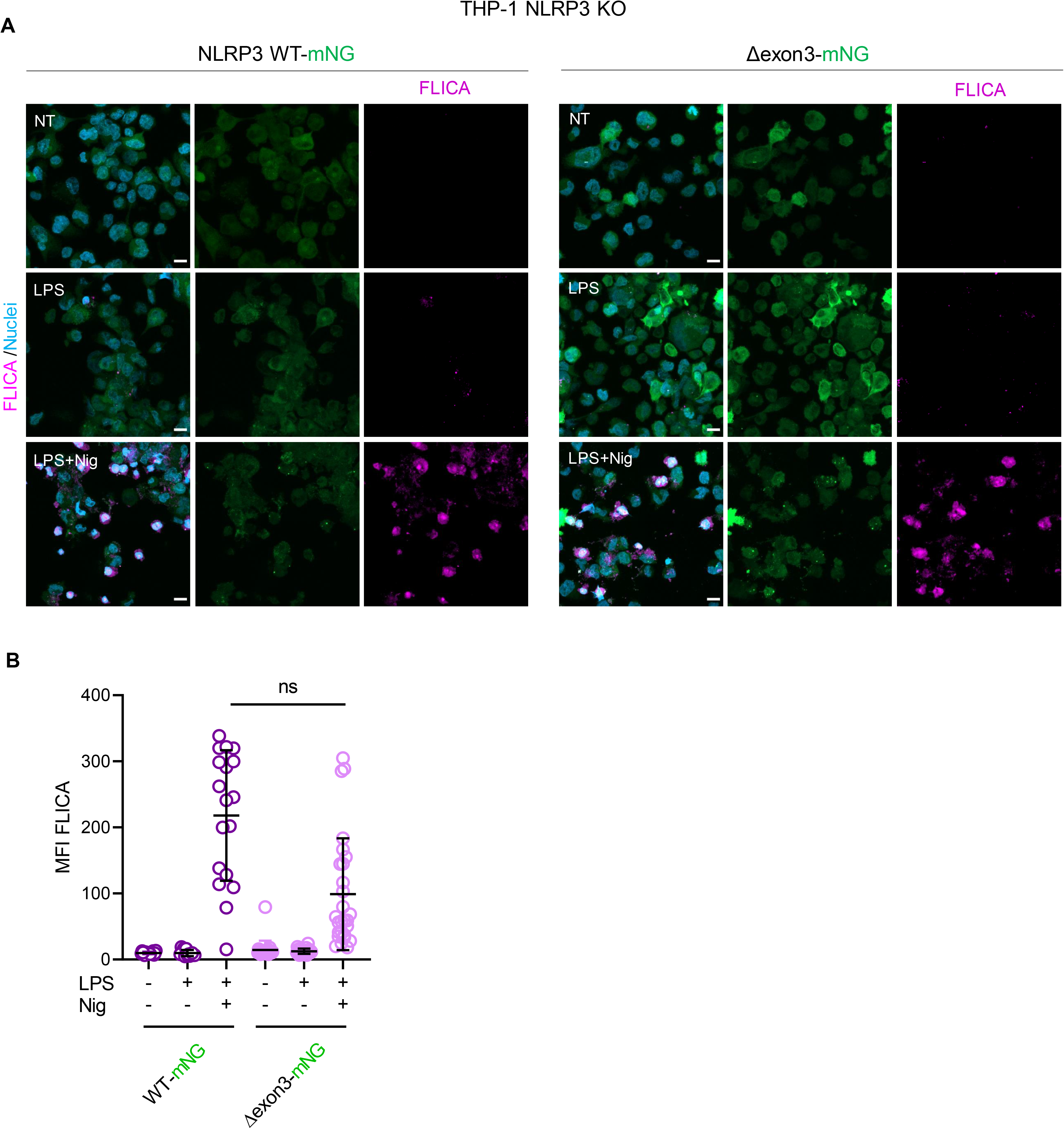
Nigericin stimulation leads to caspase-1 processing to the same extent in NLRP3 WT and Δexon3 THP-1 cells. **A)** Representative confocal microscopy images of FAM-FLICA™ substrate cleavage (magenta) by active caspase-1 in the indicated LPS-primed and nigericin-stimulated THP-1 cell lines. N=1 independent experiment. **B)** Quantification of the FLICA MFI signal in cells treated as in **A**. Each data point represents one cell from 1 independent experiment. ns=not significant according to Kruskal-Wallis test with Dunńs multiple comparisons test.

**Figure S4.**
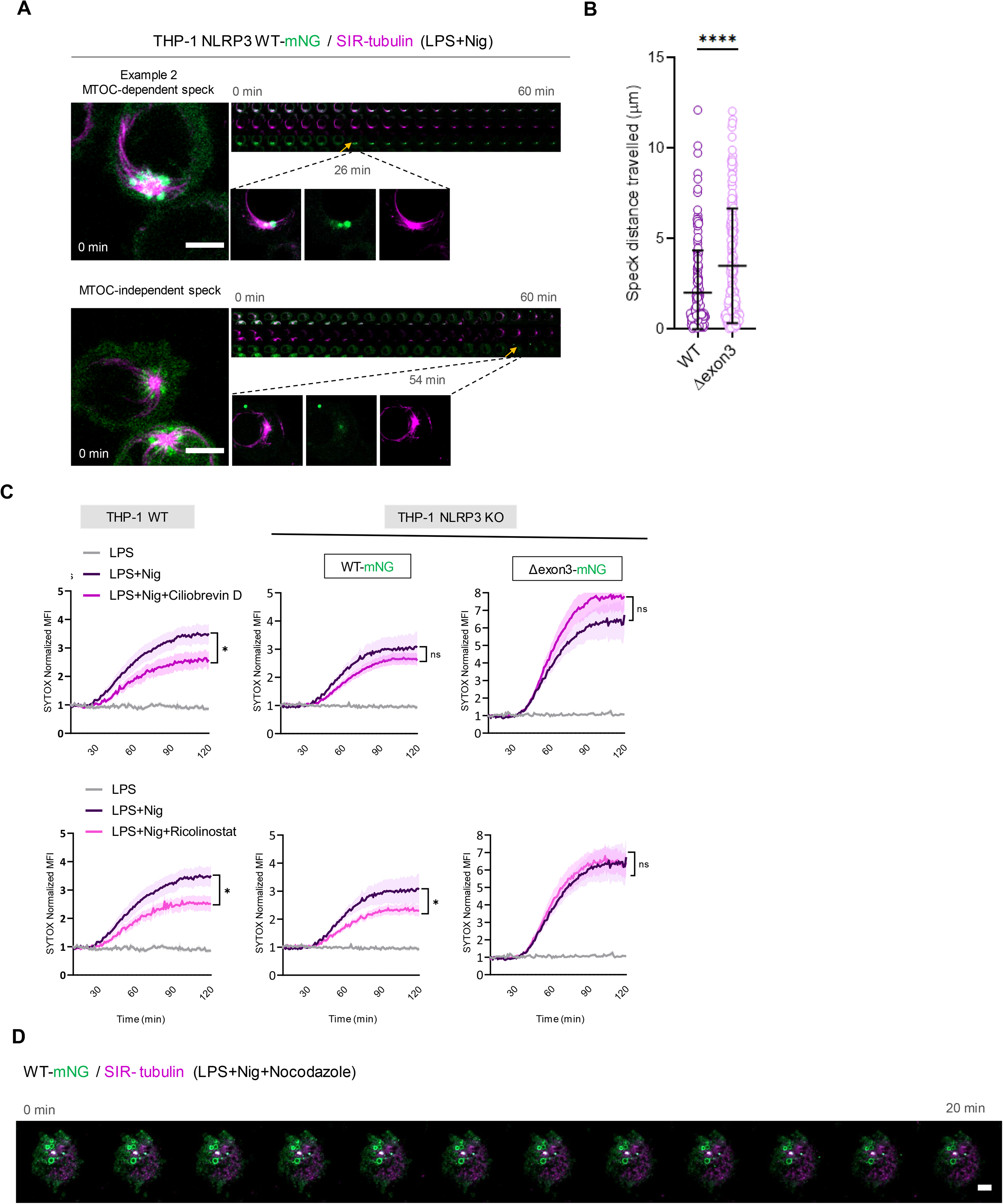
NLRP3 MTOC-independent specks are more motile and independent of microtubule trafficking inhibitors. **A)** Representative live cell imaging of the MTOC-independent NLRP3 speck formation from LPS-primed and nigericin-stimulated THP-1 NLRP3 WT-mNG cells. Microtubules were stained with the specific probe SIR-tubulin (magenta). Sequential images acquired every 3 min. Scale bar 5 µm. N=3 independent experiments. **B)** Travelled distance (μm) by WT or Δexon3 specks in cells using Mtrack2 plugin from Fiji. Each data point represents one speck. N=3 independent experiments (combined). **C)** Cell death in time measured by SYTOX orange uptake from the indicated THP-1 cell lines upon ciliobrevin (dynein inhibitor) and ricolinostat (HDAC6 inhibitor). N=3 independent experiments. *p≤ 0.05; ns, non-significant according to one-way ANOVA. **D)** Representative live cell imaging of NLRP3 WT-mNG THP-1 cells pretreated with nocodazole and then nigericin-stimulated. Sequential images acquired every 75 s. N=3 independent experiments.

**Figure S5.**
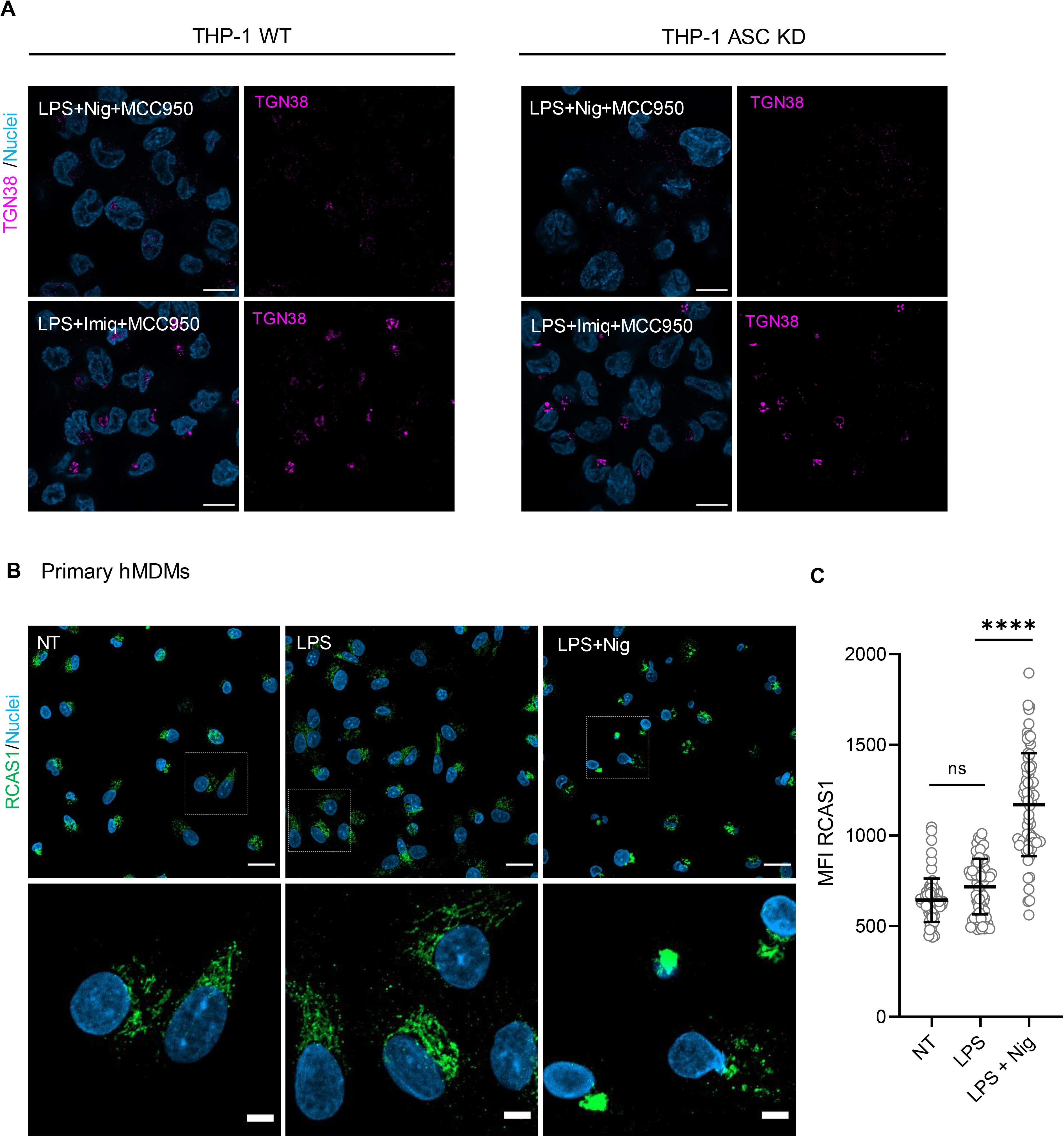
TGN aggregation is a feature of NLRP3 activation and is ASC-dependent. **A)** Representative immunofluorescence micrographs of THP-1 WT or *ASC* KD cell lines stimulated with nigericin or imiquimod in the presence of the NLRP3-specific inhibitor MCC950. TGN marked with the TGN38 antibody (magenta) and nuclei with Hoechst 33342 (blue). Scale bar 10 µm. N=3 independent experiments. **B)** Representative immunofluorescence images of LPS-primed and nigericin-stimulated human primary macrophages. Golgi membranes are marked with the RCAS antibody (green) and nuclei with Hoechst 33342 (blue). Scale bar 20 µm, zoom in 5 µm. N=3 independent experiments. **C)** Quantification of the Golgi aggregation of **B** (measured by the RCAS mean fluorescence intensity, MFI). Each data point represents one cell from 2 independent experiments. ****p≥0.0001, ns=not significant; according to Kruskal-Wallis test with Dunńs multiple comparisons test.

**Supplementary movie 1.** Live cell imaging of LPS-primed THP-1 *NLRP3* KO cell line reconstituted with WT NLRP3 forming an NLRP3 MTOC-dependent speck upon nigericin stimulation. NLRP3 (green) and microtubules (magenta).

**Supplementary movie 2.** Live cell imaging of LPS-primed THP-1 *NLRP3* KO cell line reconstituted with Δexon3 NLRP3 forming an NLRP3 MTOC-independent speck upon nigericin stimulation. NLRP3 (green) and microtubules (magenta).

**Supplementary movie 3.** As in movie 1 but showing the formation of an WT NLRP3-MTOC independent speck.

**Supplementary movie 4.** As in movie 1 but showing the formation of an WT NLRP3-MTOC independent speck in the presence of nocodazole.

**Supplementary Table 1.** Antibodies used for immunoblotting and immunofluorescence

| Antibodies | Name | Use | Company/Catalog number |
| --- | --- | --- | --- |
| α-NLRP3 | NLRP3 (D4D8T) rabbit mAb | IB: 1:1000 | Cell Signaling Technology, 15101 |
| α-NLRP3 | NLRP3 (Cryo2) mouse mAb | IB: 1:1000<br>IF: 1:100 | Adipogen, AG-20B-0014-C100 |
| α-ASC | ASC mouse mAb | IB: 1:1000<br>IF: 1:100 | Santa Cruz Biotech, sc271054 |
| α-NEK7 | NEK7 rabbit mAb | IB: 1:1000 | Cell Signaling Technology, C34C3 |
| α-Caspase-1 | Caspase-1 (D7F10) Rabbit mAb | IB: 1:1000 | Cell Signaling Technology, 3866S |
| α-IL-1β | IL-1β Mouse | IB: 1:1000 | R&D Systems |
| GSDMD | GSDMD rabbit monoclonal | IB: 1:1000 | Cell Signaling Technology (E951X) Cat. 39754 |
| α-TGN38 | TGN38 (2F 7.1) mouse mAb | IB: 1:500 | Novus Biological, NB300-575 |
| α-TGN38 | TGN38 rabbit mAb | IF: 1:100 | Novus Biological, NBP1-03495SS |
| α-RCAS1 | RCAS1 (D2B6N) rabbit mAb | IF: 1:100 | Cell signaling 12290S |
| EEA1 | EEA1 mouse monoclonal | IB: 1:1000 | BD Biosciences |
| α-γ-Tubulin | γ-Tubulin mouse mAb | IB: 1:2000<br>IF: 1:100 | Sigma T5326 |
| α-TOMM20 | TOMM20 rabbit polyclonal | IB: 1:1000 | Novus NBP1-81556 |
| α-β actin | β-actin mouse monoclonal | IB: 1:2000 | Sigma A5441-100UL |
| α-Rabbit IgG-HRP | Goat Anti-Rabbit IgG Antibody (H+L), Peroxidase | IB: 1:5000<br>or 1:10000 | Vector laboratories Inc./PI-1000 |
| α-Mouse IgG-HRP | Anti-Mouse IgG (H+L), HRP Conjugate | IB: 1:5000<br>or 1:10000 | Promega/W402B |
| α-Rabbit IgG<br>Alexa 647 | Alexa Fluor™ 647 Goat anti-Rabbit IgG (H+L) | IF: 1:500 | Invitrogen/ A-21245 |
| α-Rabbit IgG<br>Alexa 488 | Alexa Fluor™ 488 Goat anti-Rabbit IgG (H+L) | IF: 1:500 | Invitrogen/A-11008 |
| α-Mouse IgG<br>Alexa 647 | Alexa Fluor™ 647 Chicken anti-Mouse IgG (H+L) | IF: 1:500 | Invitrogen/ A-21463 |

## References

1. Schroder, K., and Tschopp, J. (2010). The inflammasomes. Cell 140, 821–832. 10.1016/J.CELL.2010.01.040.

2. Swanson, K. V., Deng, M., and Ting, J.P.Y. (2019). The NLRP3 inflammasome: molecular activation and regulation to therapeutics. Nat. Rev. Immunol. 19, 477–489. 10.1038/s41577-019-0165-0.

3. Weber, A.N.R., Tapia-Abellán, A., Liu, X., Dickhöfer, S., Aróstegui, J.I., Pelegrín, P., Welzel, T., and Kuemmerle-Deschner, J.B. (2022). Effective ex vivo inhibition of cryopyrin-associated periodic syndrome (CAPS)-associated mutant NLRP3 inflammasome by MCC950/CRID3. Rheumatology 61, e299–e313. 10.1093/rheumatology/keac273.

4. Mangan, M.S.J., Olhava, E.J., Roush, W.R., Seidel, H.M., Glick, G.D., and Latz, E. (2018). Targeting the NLRP3 inflammasome in inflammatory diseases. Nat. Rev. Drug Discov. 10.1038/nrd.2018.97.

5. Weber, A.N.R., Bittner, Z.A., Shankar, S., Liu, X., Chang, T.-H., Jin, T., and Tapia-Abellán, A. (2020). Recent insights into the regulatory networks of NLRP3 inflammasome activation. J. Cell Sci. 133. 10.1242/jcs.248344.

6. Seoane, P.I., Lee, B., Hoyle, C., Yu, S., Lopez-Castejon, G., Lowe, M., and Brough, D. (2020). The NLRP3–inflammasome as a sensor of organelle dysfunction. J. Cell Biol. 219. 10.1083/jcb.202006194.

7. Muñoz-Planillo, R., Kuffa, P., Martínez-Colón, G., Smith, B.L., Rajendiran, T.M., and Núñez, G. (2013). K+ Efflux Is the Common Trigger of NLRP3 Inflammasome Activation by Bacterial Toxins and Particulate Matter. Immunity 38, 1142–1153. 10.1016/j.immuni.2013.05.016.

8. Groß, C.J., Mishra, R., Schneider, K.S., Médard, G., Wettmarshausen, J., Dittlein, D.C., Shi, H., Gorka, O., Koenig, P.-A., Fromm, S., et al. (2016). K + Efflux-Independent NLRP3 Inflammasome Activation by Small Molecules Targeting Mitochondria. Immunity 45, 761–773. 10.1016/j.immuni.2016.08.010.

9. Tapia-Abellán, A., Angosto-Bazarra, D., Martínez-Banaclocha, H., de Torre-Minguela, C., Cerón-Carrasco, J.P., Pérez-Sánchez, H., Arostegui, J.I., and Pelegrin, P. (2019). MCC950 closes the active conformation of NLRP3 to an inactive state. Nat. Chem. Biol. 15, 560–564. 10.1038/s41589-019-0278-6.

10. Tapia-Abellán, A., Angosto-Bazarra, D., Alarcón-Vila, C., Baños, M.C., Hafner-Bratkovič, I., Oliva, B., and Pelegrín, P. (2021). Sensing low intracellular potassium by NLRP3 results in a stable open structure that promotes inflammasome activation. Sci. Adv. 7, 4468–4483. 10.1126/sciadv.abf4468.

11. He, W., Wan, H., Hu, L., Chen, P., Wang, X., Huang, Z., Yang, Z.-H., Zhong, C.-Q., and Han, J. (2015). Gasdermin D is an executor of pyroptosis and required for interleukin-1β secretion. Cell Res. 25, 1285– 1298. 10.1038/cr.2015.139.

12. Akbal, A., Dernst, A., Lovotti, M., Mangan, M.S.J., McManus, R.M., and Latz, E. (2022). How location and cellular signaling combine to activate the NLRP3 inflammasome. Cell. Mol. Immunol. 19, 1201–1214. 10.1038/s41423-022-00922-w.

13. Andreeva, L., David, L., Rawson, S., Shen, C., Pasricha, T., Pelegrin, P., and Wu, H. (2021). NLRP3 cages revealed by full-length mouse NLRP3 structure control pathway activation. Cell 184, 6299–6312.e22. 10.1016/j.cell.2021.11.011.

14. Hochheiser, I. V., Pilsl, M., Hagelueken, G., Moecking, J., Marleaux, M., Brinkschulte, R., Latz, E., Engel, C., and Geyer, M. (2022). Structure of the NLRP3 decamer bound to the cytokine release inhibitor CRID3. Nature 604, 184–189. 10.1038/s41586-022-04467-w.

15. Hu, Z., Yan, C., Liu, P., Huang, Z., Ma, R., Zhang, C., Wang, R., Zhang, Y., Martinon, F., Miao, D., et al. (2013). Crystal structure of NLRC4 reveals its autoinhibition mechanism. Science 341, 172–175. 10.1126/SCIENCE.1236381.

16. Zhang, L., Chen, S., Ruan, J., Wu, J., Tong, A.B., Yin, Q., Li, Y., David, L., Lu, A., Wang, W.L., et al. (2015). Cryo-EM structure of the activated NAIP2-NLRC4 inflammasome reveals nucleated polymerization. Science (80-.). 350, 404–409. 10.1126/science.aac5789.

17. Sharif, H., Wang, L., Wang, W.L., Magupalli, V.G., Andreeva, L., Qiao, Q., Hauenstein, A. V., Wu, Z., Núñez, G., Mao, Y., et al. (2019). Structural mechanism for NEK7-licensed activation of NLRP3 inflammasome. Nature. 10.1038/s41586-019-1295-z.

18. Dekker, C., Mattes, H., Wright, M., Boettcher, A., Hinniger, A., Hughes, N., Kapps-Fouthier, S., Eder, J., Erbel, P., Stiefl, N., et al. (2021). Crystal Structure of NLRP3 NACHT Domain With an Inhibitor Defines Mechanism of Inflammasome Inhibition. J. Mol. Biol. 433, 167309. 10.1016/j.jmb.2021.167309.

19. Hafner-Bratkovič, I., Sušjan, P., Lainšček, D., Tapia-Abellán, A., Cerović, K., Kadunc, L., Angosto-Bazarra, D., Pelegrin, P., and Jerala, R. (2018). NLRP3 lacking the leucine-rich repeat domain can be fully activated via the canonical inflammasome pathway. Nat. Commun. 9, 5182. 10.1038/s41467-018-07573-4.

20. Chen, J., and Chen, Z.J. (2018). PtdIns4P on dispersed trans-Golgi network mediates NLRP3 inflammasome activation. Nature 564, 71–76. 10.1038/s41586-018-0761-3.

21. Zhang, Z., Meszaros, G., He, W., Xu, Y., de Fatima Magliarelli, H., Mailly, L., Mihlan, M., Liu, Y., Puig Gámez, M., Goginashvili, A., et al. (2017). Protein kinase D at the Golgi controls NLRP3 inflammasome activation. J. Exp. Med. 214, 2671–2693. 10.1084/jem.20162040.

22. Zhang, Z., Venditti, R., Ran, L., Liu, Z., Vivot, K., Schürmann, A., Bonifacino, J.S., De Matteis, M.A., and Ricci, R. (2023). Distinct changes in endosomal composition promote NLRP3 inflammasome activation. Nat. Immunol. 24, 30–41. 10.1038/s41590-022-01355-3.

23. Magupalli, V.G., Negro, R., Tian, Y., Hauenstein, A. V., Di Caprio, G., Skillern, W., Deng, Q., Orning, P., Alam, H.B., Maliga, Z., et al. (2020). HDAC6 mediates an aggresome-like mechanism for NLRP3 and pyrin inflammasome activation. Science (80-.). 369. 10.1126/science.aas8995.

24. Ohto, U., Kamitsukasa, Y., Ishida, H., Zhang, Z., Murakami, K., Hirama, C., Maekawa, S., and Shimizu, T. (2022). Structural basis for the oligomerization-mediated regulation of NLRP3 inflammasome activation. Proc. Natl. Acad. Sci. 119. 10.1073/pnas.2121353119.

25. Xiao, L., Magupalli, V.G., and Wu, H. (2023). Cryo-EM structures of the active NLRP3 inflammasome disc. Nature 613, 595–600. 10.1038/s41586-022-05570-8.

26. Bittner, Z.A., Liu, X., Mateo Tortola, M., Tapia-Abellán, A., Shankar, S., Andreeva, L., Mangan, M., Spalinger, M., Kalbacher, H., Düwell, P., et al. (2021). BTK operates a phospho-tyrosine switch to regulate NLRP3 inflammasome activity. J. Exp. Med. 218, 19. 10.1084/jem.20201656.

27. Simon, M.L.A., Platre, M.P., Assil, S., van Wijk, R., Chen, W.Y., Chory, J., Dreux, M., Munnik, T., and Jaillais, Y. (2014). A multi-colour/multi-affinity marker set to visualize phosphoinositide dynamics in Arabidopsis. Plant J. 77, 322–337. 10.1111/tpj.12358.

28. Omrane, M., Camara, A.S., Taveneau, C., Benzoubir, N., Tubiana, T., Yu, J., Guérois, R., Samuel, D., Goud, B., Poüs, C., et al. (2019). Septin 9 has Two Polybasic Domains Critical to Septin Filament Assembly and Golgi Integrity. iScience 13, 138–153. 10.1016/j.isci.2019.02.015.

29. Hoss, F., Mueller, J.L., Rojas Ringeling, F., Rodriguez-Alcazar, J.F., Brinkschulte, R., Seifert, G., Stahl, R., Broderick, L., Putnam, C.D., Kolodner, R.D., et al. (2019). Alternative splicing regulates stochastic NLRP3 activity. Nat. Commun. 10, 3238. 10.1038/s41467-019-11076-1.

30. Zheng, Y., Jung, M.K., and Oakley, B.R. (1991). γ-Tubulin is present in Drosophila melanogaster and homo sapiens and is associated with the centrosome. Cell 65, 817–823. 10.1016/0092-8674(91)90389-G.

31. Lukinavičius, G., Reymond, L., D’Este, E., Masharina, A., Göttfert, F., Ta, H., Güther, A., Fournier, M., Rizzo, S., Waldmann, H., et al. (2014). Fluorogenic probes for live-cell imaging of the cytoskeleton. Nat. Methods 11, 731–733. 10.1038/nmeth.2972.

32. Martinon, F., Pétrilli, V., Mayor, A., Tardivel, A., and Tschopp, J. (2006). Gout-associated uric acid crystals activate the NALP3 inflammasome. Nature 440, 237–241. 10.1038/nature04516.

33. Liu, X., Pichulik, T., Wolz, O.-O., Dang, T.-M., Stutz, A., Dillen, C., Delmiro Garcia, M., Kraus, H., Dickhöfer, S., Daiber, E., et al. (2017). Human NACHT, LRR, and PYD domain–containing protein 3 (NLRP3) inflammasome activity is regulated by and potentially targetable through Bruton tyrosine kinase. J. Allergy Clin. Immunol. 140, 1054–1067.e10. 10.1016/j.jaci.2017.01.017.

34. Schmacke, N.A., O’Duill, F., Gaidt, M.M., Szymanska, I., Kamper, J.M., Schmid-Burgk, J.L., Mädler, S.C., Mackens-Kiani, T., Kozaki, T., Chauhan, D., et al. (2022). IKKβ primes inflammasome formation by recruiting NLRP3 to the trans-Golgi network. Immunity 55, 2271–2284.e7. 10.1016/j.immuni.2022.10.021.

35. Lee, B., Hoyle, C., Wellens, R., Green, J.P., Martin-Sanchez, F., Williams, D.M., Matchett, B.J., Seoane, P.I., Bennett, H., Adamson, A., et al. (2023). Disruptions in endocytic traffic contribute to the activation of the NLRP3 inflammasome. Sci. Signal. 16, eabm7134. 10.1126/scisignal.abm7134.

36. Machtens, D.A., Bresch, I.P., Eberhage, J., Reubold, T.F., and Eschenburg, S. (2022). The Inflammasome Activity of NLRP3 Is Independent of NEK7 in HEK293 Cells Co-Expressing ASC. Int. J. Mol. Sci. 23, 10269. 10.3390/ijms231810269.

37. Siddhanta, A., Backer, J.M., and Shields, D. (2000). Inhibition of Phosphatidic Acid Synthesis Alters the Structure of the Golgi Apparatus and Inhibits Secretion in Endocrine Cells. J. Biol. Chem. 275, 12023– 12031. 10.1074/jbc.275.16.12023.

38. Thakur, R., Naik, A., Panda, A., and Raghu, P. (2019). Regulation of Membrane Turnover by Phosphatidic Acid: Cellular Functions and Disease Implications. Front. Cell Dev. Biol. 7. 10.3389/fcell.2019.00083.

39. Baron, C.L., and Malhotra, V. (2002). Role of Diacylglycerol in PKD Recruitment to the TGN and Protein Transport to the Plasma Membrane. Science (80-.). 295, 325–328. 10.1126/science.1066759.

40. Freyberg, Z. (2003). “Slip, sliding away”: phospholipase D and the Golgi apparatus. Trends Cell Biol. 13, 540–546. 10.1016/j.tcb.2003.08.004.

41. Marat, A.L., and Haucke, V. (2016). Phosphatidylinositol 3-phosphates—at the interface between cell signalling and membrane traffic. EMBO J. 35, 561–579. 10.15252/embj.201593564.

42. Hornung, V., Bauernfeind, F., Halle, A., Samstad, E.O., Kono, H., Rock, K.L., Fitzgerald, K.A., and Latz, E. (2008). Silica crystals and aluminum salts activate the NALP3 inflammasome through phagosomal destabilization. Nat. Immunol. 9, 847–856. 10.1038/ni.1631.

43. Karasawa, T., Komada, T., Yamada, N., Aizawa, E., Mizushina, Y., Watanabe, S., Baatarjav, C., Matsumura, T., and Takahashi, M. (2022). Cryo-sensitive aggregation triggers NLRP3 inflammasome assembly in cryopyrin-associated periodic syndrome. Elife 11, 1–26. 10.7554/eLife.75166.

44. Slobodnick, A., Shah, B., Pillinger, M.H., and Krasnokutsky, S. (2015). Colchicine: Old and New. Am. J. Med. 128, 461–470. 10.1016/j.amjmed.2014.12.010.

45. Kuemmerle-Deschner, J.B., Gautam, R., George, A.T., Raza, S., Lomax, K.G., and Hur, P. (2020). Systematic literature review of efficacy/effectiveness and safety of current therapies for the treatment of cryopyrin-associated periodic syndrome, hyperimmunoglobulin D syndrome and tumour necrosis factor receptor-associated periodic syndrome. RMD Open 6, e001227. 10.1136/rmdopen-2020-001227.

46. Campeau, E., Ruhl, V.E., Rodier, F., Smith, C.L., and Rahmberg, B.L. (2009). A Versatile Viral System for Expression and Depletion of Proteins in Mammalian Cells. PLoS One 4, 6529. 10.1371/journal.pone.0006529.

47. Schink, K.O., Tan, K.W., Spangenberg, H., Martorana, D., Sneeggen, M., Stévenin, V., Enninga, J., Campsteijn, C., Raiborg, C., and Stenmark, H. (2021). The phosphoinositide coincidence detector Phafin2 promotes macropinocytosis by coordinating actin organisation at forming macropinosomes. Nat. Commun. 12. 10.1038/s41467-021-26775-x.

48. Nguyen, Q., and Yokota, T. (2019). Antisense oligonucleotides for the treatment of cardiomyopathy in Duchenne muscular dystrophy. Am J Transl Res 11, 1202–1218.

49. Desmet, F.O., Hamroun, D., Lalande, M., Collod-Bëroud, G., Claustres, M., and Béroud, C. (2009). Human Splicing Finder: An online bioinformatics tool to predict splicing signals. Nucleic Acids Res. 37, 1–14. 10.1093/nar/gkp215.

50. Zuker, M. (2003). Mfold web server for nucleic acid folding and hybridization prediction. Nucleic Acids Res. 31, 3406–3415. 10.1093/nar/gkg595.

